# Diverse spatial organization of photosynthetic membranes among purple nonsulfur bacteria

**DOI:** 10.1101/248997

**Authors:** Breah LaSarre, David T. Kysela, Barry D. Stein, Adrien Ducret, Yves V. Brun, James B. McKinlay

## Abstract

In diverse bacteria, proper cellular physiology requires the utilization of protein- or membrane-bound compartments that afford specific metabolic capabilities. One such compartment is the light-harvesting intracytoplasmic membrane (ICM) of purple nonsulfur bacteria (PNSB). Here we reveal that ICMs are subject to differential spatial organization among PNSB. We visualized ICMs in live cells of fourteen PNSB species by exploiting the natural autofluorescence of the photosynthetic machinery. We then quantitatively characterized ICM localization using automated computational analysis of autofluorescence patterns within single cells across the population. Our studies revealed that ICMs are localized in distinct subcellular patterns that differ between species; some PNSB elaborate ICMs throughout the cell, while others spatially restrict ICM to varying degrees. The most highly-restricted ICMs were localized in a specific pattern corresponding to progression of cell growth and division. An identical pattern of ICM restriction was conserved across at least two genera. Phylogenetic and phenotypic comparisons established that ICM localization and ICM architecture are not strictly interdependent and that neither trait fully correlates with the evolutionary relatedness of the species. This discovery of new diversity in bacterial cell organization has implications for understanding both the mechanisms underpinning spatial arrangement of bacterial compartments and the potential benefits of adopting different spatiotemporal patterns.

## INTRODUCTION

In bacteria and eukaryotes alike, proper cellular physiology relies on robust subcellular organization. Diverse bacteria organize their cytoplasmic milieu by constructing specialized protein- and membrane-bound compartments^1^. Among these compartments are the intracellular membranes (ICMs) of purple nonsulfur bacteria (PNSB). ICMs, also known as chromatophores, house the proteins and pigments responsible for photosynthesis^2^, akin to the thylakoid membranes of cyanobacteria and plant chloroplasts. ICMs originate from invaginations of the cytoplasmic membrane (CM)^3^, remain physically contiguous with the CM following their elaboration^4-7^, and are presumed to enhance the efficiency of light capture and energy transformation^5,8,9^. However, amid their comparable origin and function, ICMs exhibit species-specific architectures, ranging from vesicles to tubes to lamellae^3^ (Fig. 1). ICM architectures are currently understood to result from inter-molecular interactions between components of the photosynthetic machinery^10-13^.

**Fig. 1.**
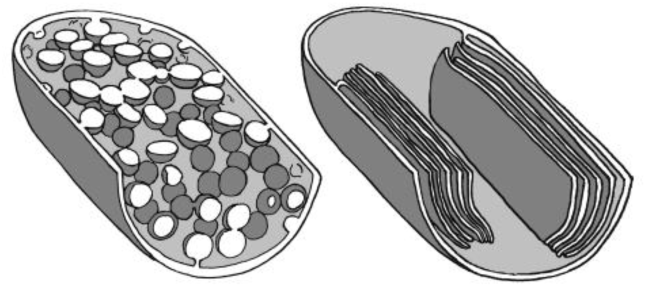
Illustration of vesicular (left) and lamellar (right) ICM architectures. Quarter-cell illustrations (not to scale) are based on published EM images of *Rba. sphaeroides* and *Rps. palustris*, respectively.

Most PNSB synthesize ICMs only under low-oxygen conditions that allow anoxygenic photosynthesis^14-16^; cells grown anaerobically in light (phototrophically) contain ICMs, whereas those grown aerobically in darkness (chemotrophically) do not. ICMs have been used as model systems for studying membrane biogenesis and photochemistry^2,3,17^, but remain poorly characterized with regard to cellular organization, presumably because studies relied on biochemical analysis of bulk ICM and electron microscopy of a small number of cells. Furthermore, while PNSB comprise more than 20 genera^2^, most ICM studies have focused on only a handful of species of *Rhodobacter, Rhodospirillum*, and *Rhodopseudomonas*. PNSB are phylogenetically and physiologically diverse^2^, thus it would be premature to assume that all species coordinate ICMs equivalently. In line with this notion, historical observations have alluded to differential ICM organization between species; *Rhodopseudomonas palustris* might restrict ICMs to regions of the cell^18^, while other PNSB, such as *Rhodobacter sphaeroides* and *Rhodospirillum rubrum*, elaborate ICMs throughout the cell^4,10^. Bearing this in mind, we sought to determine whether ICMs are subject to spatial organization and, if so, whether such organization is conserved or variable among PNSB.

## RESULTS

### Bacteriochlorophyll autofluorescence is a non-invasive tool for visualizing ICMs in live cells

We first set out to develop a method for visualizing the ICM-residing photosynthetic machinery (photosystems) using the natural fluorescence (autofluorescence) of bacteriochlorophyll (BChl); this approach emulates the use of chlorophyll autofluorescence to visualize photosynthetic components within plant chloroplasts and cyanobacteria^19,20^. We deemed BChl to be a suitable ICM marker for several reasons. First, BChl only accumulates under conditions that stimulate ICM and photosystem synthesis^16,21^. Second, BChl is an obligate component of PNSB photosystems and is thus essential for ICM synthesis^3^. Finally, BChl exists almost entirely within photosystems, with very little free in the cytoplasm^22^, and these photosystems predominantly reside within ICMs^7,15,23,24^.

We tested the hypothesis that BChl autofluorescence could be used to visualize ICMs using the model PNSB species, *Rps. palustris*. In accord with oxygen-regulated ICM synthesis^16^, aerobically-grown cells lacking ICMs were deficient for pigmentation, whereas phototrophically-grown cells containing ICM were pigmented and exhibited characteristic photosystem-associated absorbance peaks (Fig. 2a,b). When we examined *Rps. palustris* by epifluorescence microscopy, phototrophic cells exhibited autofluorescence detected using DAPI filter sets (Fig. 2c, Supplementary Fig. 1). No autofluorescence was detected using several other common filter sets (Fig. 2c). In contrast to phototrophic cells, there was no autofluorescence detected in aerobic cells using any of these filter sets (Fig. 2c, Supplementary Fig. 1). Thus, autofluorescence was specific to ICM-containing cells.

**Fig. 2.**
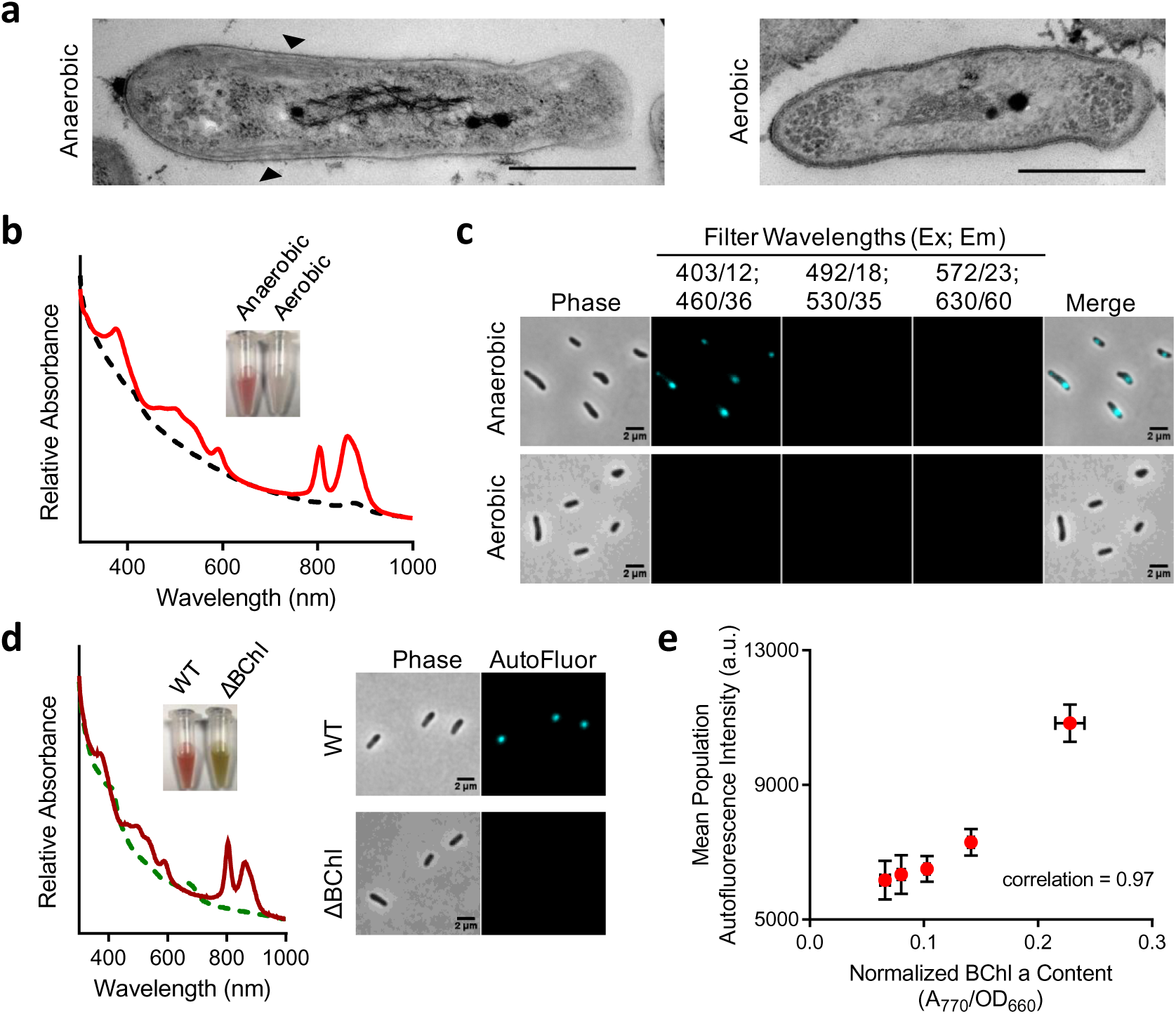
Bacteriochlorophyll autofluorescence as a proxy for ICMs in live cells. **a**, Electron microscopy images of wild-type (WT) *Rps. palustris* strain CGA009 cells grown in PMsuccYE either anaerobically in 8 μmol s^−1^ m^−2^ light (left) or aerobically in darkness (right). Scale bar, 500 nm. Arrowheads indicate ICMs. **b**, Spectral analysis of WT *Rps. palustris* cells grown anaerobically (solid red line) or aerobically (dashed black line) as described for **a**. *Inset*, photograph of concentrated cells grown under each condition. **c**, Fluorescence microscopy images of cells from cultures used in **b** using indicated filter sets (Excitation center/range; Emission center/range). Microscopy lookup tables (LUTs) for contrast and brightness are equivalent between fluorescence panels. **d**, Spectral analysis (*left*) and microscopy images (*right*) of WT (solid dark red line) and BChl-deficient ΔBChl) (dashed green line) *Rps. palustris*, each grown by anaerobic respiration with N_2_O in darkness. Autofluorescence (AutoFluor) was detected using the 403/12;460/36 filter set. LUTs are equivalent between panels. *Inset left*, photograph of concentrated cells of each strain. **e**, Correlation analysis of BChl *a* content and cellular autofluorescence intensity. Points represent mean population cellular autofluorescence intensity of independent cultures of WT *Rps. palustris* grown anaerobically in PMsuccYE in light intensities between 8-60 μmol s^−1^ m^−2^. Autofluorescence (background-corrected mean cellular intensity) was measured using MicrobeJ. Error bars, SD of 3 technical replicates performed over 2 days. For technical replicates, 8 μmol s^−1^ m^−2^, n = 251, 869, 1337 cells; 20 μmol s^−1^ m^−2^, n = 382, 930, 1535 cells; 32 μmol s^−1^ m^−2^, n = 392, 876, 1420 cells; 50 μmol s^−1^ m^−2^, n = 421, 786, 1101 cells; 60 μmol s^−1^ m^−2^, n = 300, 973, 1419 cells.

We next verified that autofluorescence was derived from BChl using a mutant lacking BChl. As BChl is essential for phototrophic growth, we exploited a growth condition in which ICMs are induced but not required^21^. Specifically, we grew *Rps. palustris* by anaerobic respiration in darkness using N_2_O as the terminal electron acceptor^25^. In these conditions, the absence of O_2_ prompts ICM synthesis but energy is generated via respiration with N_2_O, and thus growth of a BChl-deficient mutant ΔBChl) was comparable to that of wild-type (WT) (Supplementary Fig. 2). WT *Rps. palustris* cells grown by anaerobic respiration exhibited autofluorescence whereas ΔBChl cells did not (Fig. 2d). Separately, we also examined whether other photosystem-associated pigments, carotenoids, were responsible for autofluorescence by examining a Δ*crtI* mutant that produces BChl but not carotenoids (Supplementary Fig. 3)^26^. Unlike the ΔBChl mutant, Δ*crtI* mutant cells still exhibited autofluorescence, though at a lower intensity than WT cells (Supplementary Fig. 3). The decreased intensity was not surprising given that carotenoids, while not essential for ICM synthesis, contribute to normal photosystem assembly^26-29^ and thereby affect both BChl levels and the local environment of BChl, which is known to influence BChl spectral properties^8,10,27,30^. While it is possible that carotenoids contribute to autofluorescence, our data demonstrate that carotenoids are not necessary for autofluorescence. Consistent with this conclusion, autofluorescence was detectable in a range of PNSB that all have BChl type *a* but that use different carotenoids (discussed below) (Supplementary Table 1).

We also evaluated the relationship between cellular BChl levels and autofluorescence intensity. In PNSB, BChl levels directly correlate to photosynthetic machinery and ICM abundance^4,7,31,32^, and these levels are inversely proportional to light availability^16^. We grew *Rps. palustris* in different light intensities and compared BChl content to autofluorescence intensity. In agreement with our initial hypothesis, we observed a direct correlation between cellular BChl content and cellular autofluorescence intensity (Fig. 2e). Given this correlation, and in combination with the prior data, we conclude that autofluorescence is derived from BChl and infer that BChl autofluorescence can be used as a non-invasive marker for ICM localization in live PNSB.

### ICMs are spatially restricted in *Rps. palustris* in a pattern that promotes ICM inheritance upon cell division

The autofluorescence observed in *Rps. palustris* was strikingly focal in nature; specifically, ICMs were localized to the slightly wider, ovoid region near cell poles (Fig. 2c,d, Fig. 3a). Our EM images, like others previously^18^, corroborated this localization pattern, showing peripheral stacks of lamellar ICMs near cell poles and an absence of ICMs in the narrower, so-called ‘tube’ region^18^ of longer cells (Fig. 2a). We hereon refer to this localization pattern as longitudinal ICM restriction.

**Fig. 3.**
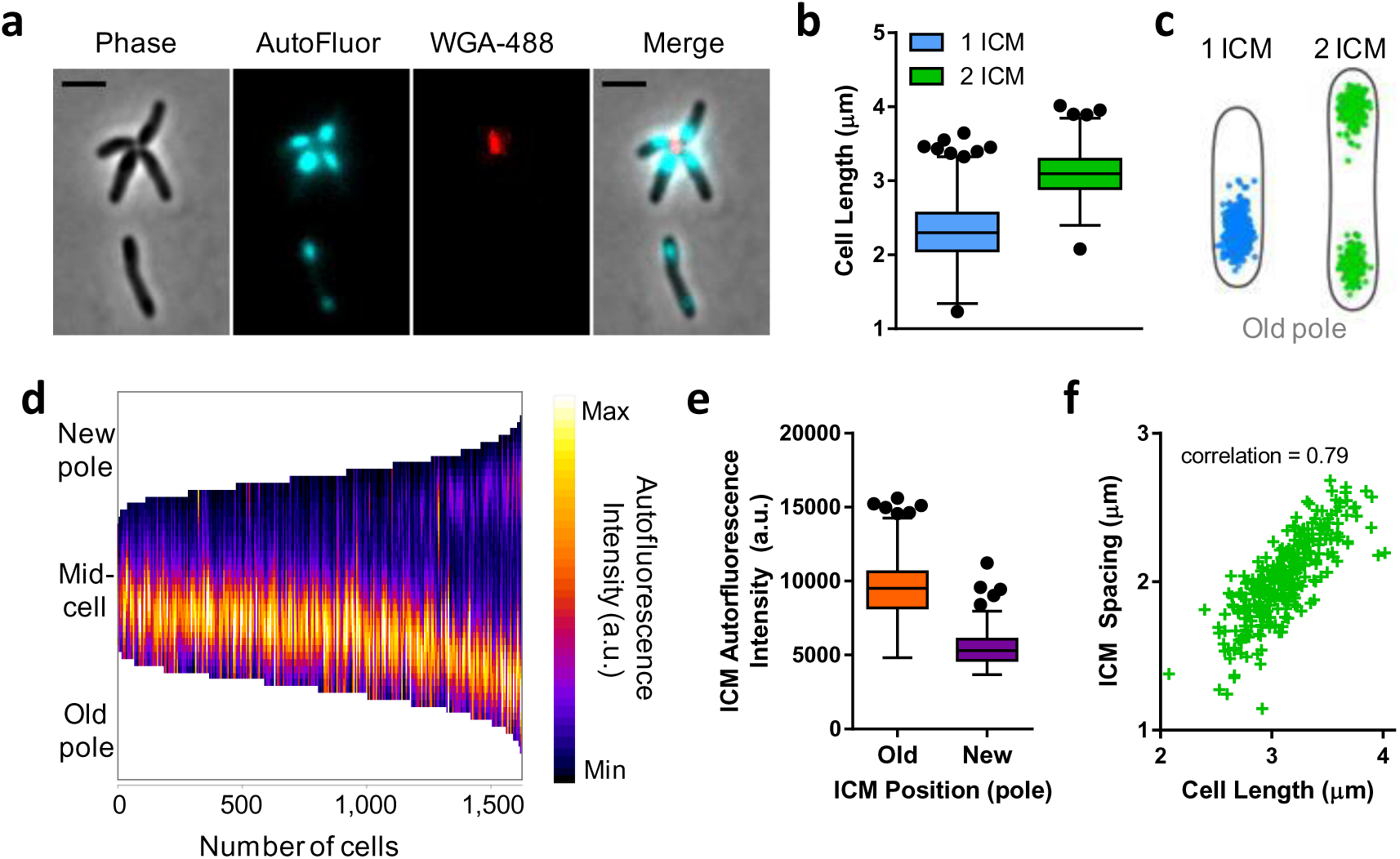
Restricted ICM localization in *Rps. palustris* is non-random. **a-f**, Analysis of ICM localization in UPP-bearing WT *Rps. palustris* cells grown in anaerobic PMsuccYE in 8 μmol s^−1^ m^−2^ light. **a**, Microscopy image of *Rps. palustris* cells showing autofluorescence (AutoFluor, cyan) and UPP stained with WGA-488 (false-colored red). Scale bar, 2 μm. **b**, Lengths of cells containing one or two ICMs. 1 ICM, n = 1330 cells; 2 ICM, n = 293 cells. **c**, Cellular position maps of autofluorescence centroids for ICMs detected in cells in b. Cell outlines depict average cell shape for each population, generated by MicrobeJ. **d**, Demograph of autofluorescence intensities measured along the medial cell axis of all cells in **b**. Cells were sorted from shortest to longest. **e**, Background-corrected mean autofluorescence intensity of ICMs located proximal (old) or distal (new) to the UPP-bearing pole in cells with 2 ICMs. n = 586 ICM maxima. **f**, Longitudinal distance (ICM spacing) between ICM centroids within cells containing two ICMs plotted as a function of cell length. n = 293 ICM pairs.

Our ability to visualize discrete ICMs prompted us to characterize the subcellular spatial distribution of ICMs across the population. As *Rps. palustris* cells divide asymmetrically and produce a unipolar polysaccharide adhesin (UPP) at the old cell pole^18,33,34^, we stained UPP with a fluorescent lectin dye to empirically orient cells (Fig. 3a). While average cell widths were comparable regardless of ICM number (data not shown), cells containing two ICMs were typically longer than cells containing a single ICM (Fig. 3b). Single ICMs were always located proximal to the UPP-bearing old pole, while in cells with two ICMs, the ICMs were positioned towards the two cell poles (Fig. 3c,d). Regardless of ICM number, the position of the UPP-proximal ICM from the UPP-bearing pole was conserved in all cells (Fig. 3c,d). A similar distance was maintained between the second ICM and the non-UPP-bearing pole in cells with two ICMs (Fig. 3c,d), although the autofluorescence intensity of the second ICM was consistently weaker than that of the first (Fig. 3d,e). As a consequence of the conserved ICM position relative to each pole, the distance between ICM pairs within single cells (ICM spacing) increased as a function of cell length (Fig. 3f). UPP is only present on a subset of cells in the population, but analysis of non-UPP-bearing cells (for which no polarity was assigned) confirmed that the longitudinally-restricted pattern of ICM localization was conserved across the population (Supplementary Fig. 4). Longitudinally-restricted ICM localization was also observed in environmental *Rps. palustris* isolates^35^ (Supplementary Fig. 5) and thus appears to be a conserved feature of this species. Additionally, longitudinal ICM restriction was maintained regardless of light availability (Supplementary Fig. 6), despite higher BChl levels and autofluorescence intensity in cells grown in lower light intensities (Fig. 1d). As such, we infer that ICM expansion necessary for accommodation of additional photosynthetic machinery occurs within resolute spatial constraints in *Rps. palustris*.

The observed correlations between cell length, ICM position, and ICM number suggested coordination of ICMs with the progression of cell growth and division. *Rps. palustris* grows polarly and divides by budding^18^. We hypothesized that single ICMs were those of mother cells and would be retained by the mother cells, while second ICMs developed near the new pole in pre-divisional cells and would be inherited by nascent daughter cells. To test this hypothesis, we monitored ICM development and position in growing *Rps. palustris* cells by time-lapse microscopy. Indeed, mother cells contained a single ICM, elongated from the opposite (new) pole, developed a second ICM of initially weaker autofluorescence near the new pole, and then completed cell division between the two ICMs (Fig. 4). Consequently, the mother and daughter cell each contained a single ICM following cell division.

**Fig. 4.**
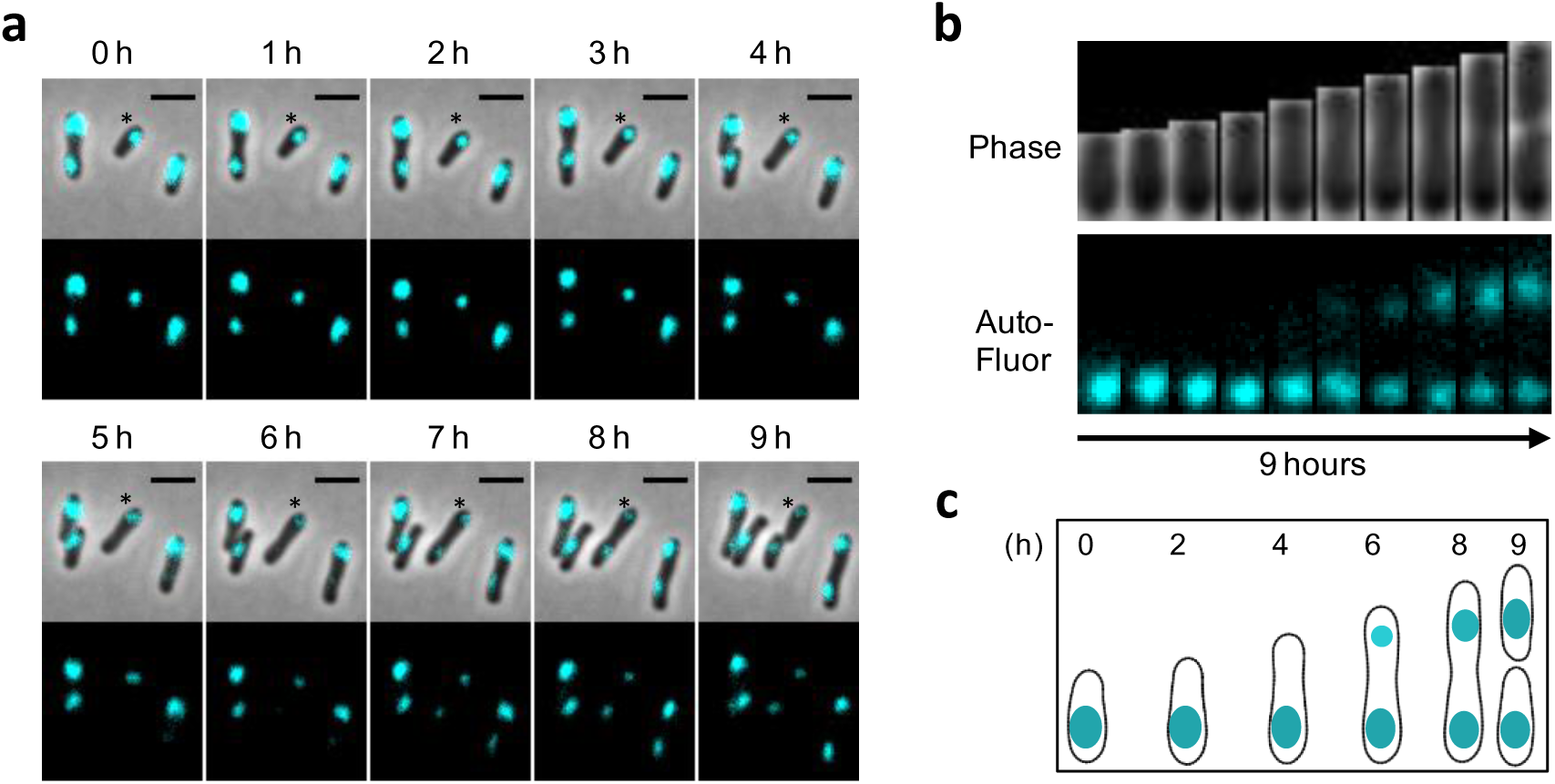
Time-lapse imaging of ICMs in *Rps. palustris*. **a**, Time-lapse merge (upper) and autofluorescence (lower) montage of *Rps. palustris* CGA009. Time elapsed is indicated above each frame. Scale bar, 2 μm. Asterisk indicates single cell depicted in subsequent panels. **b**, Phase and autofluorescence profiles of single cell from **a** over time. **c**, Schematic of cell shape (generated by MicrobeJ) and associated ICMs at indicated time points in cell in **b**.

### ICM localization and ICM architecture are not strictly interdependent, and neither trait fully correlates with species phylogeny

As ICM architectures vary between species^3^, we sought to determine whether other PNSB exhibited restricted ICM localization. We obtained 13 additional BChl type a-containing PNSB species that utilize either lamellar ICMs, similar to *Rps. palustris*, or vesicular ICMs, an architecture that has been most studied in other PNSB (Fig. 5a, Supplementary Table 1), and assessed ICM localization using BChl autofluorescence (Fig. 5b, Supplementary Fig. 7). To gain insight into the basis of ICM diversity, we mapped ICM phenotypes (localization and/or architecture) on to a phylogenetic tree that incorporated the 21 PNSB for which genome sequences are available, including six we sequenced for this study, and ICM architecture has been documented (Fig. 4, Supplementary Table 1).

**Fig. 5.**
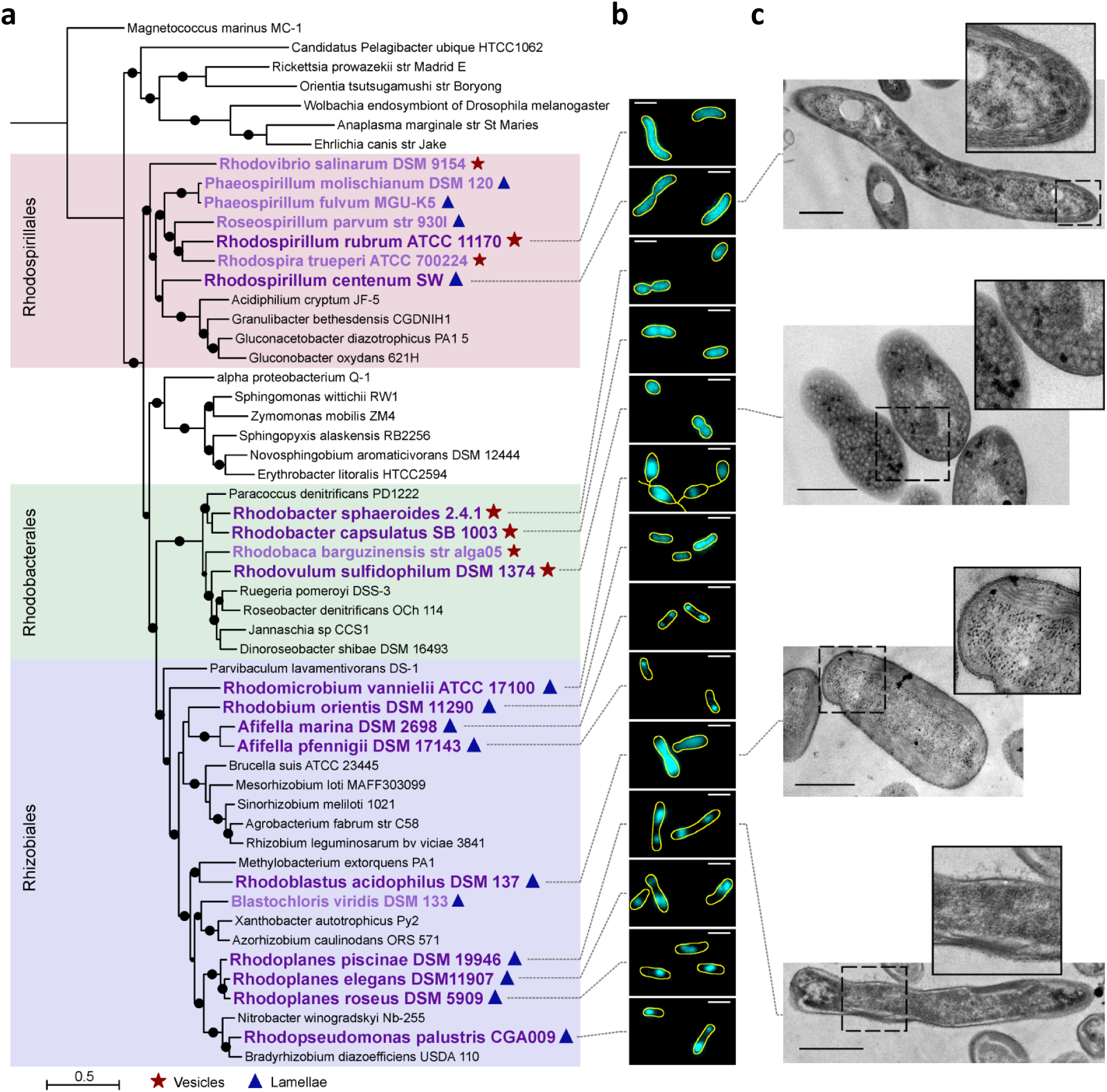
Phylogenetic analysis, autofluorescence patterns, and electron microscopy (EM) of select PNSB. **a**, Maximum-likelihood tree of non-PNSB (black text) and PNSB species (purple text) generated using a concatenated alignment of 36 housekeeping genes. *M. marinus* is included as an outgroup. Bootstrap values above 70% are indicated with circles. Historically noted ICM architectures are indicated with symbols (see legend). **b**, Autofluorescence (cyan) of select PNSB (large, dark purple text in **a**) with overlaid cell outlines (yellow, generated by FIJI). Scale bar, 2 μm. LUTs are not equivalent between images. Phase, autofluorescence and merged images are available in Supplementary Fig. 7. **c**, EM images of (top to bottom) *Rsp. centenum*, *Rvu. sulphidophilum*, *Rbl. acidophilus*, and *Rpl. piscinae*, with accompanying zoom of region indicated by dashed box for each image. Scale bar, 500 nm.

Of the 13 additional species examined by microscopy, seven (*Rsp. rubrum*, *Rhodospirillum centenum*, *Rba. sphaeroides*, *Rhodobacter capsulatus*, *Rhodovulum sulfidophilum*, *Rhodobium orientis*, and *Rhodoblastus acidophilus*) exhibited relatively uniform autofluorescence that spanned most or all of the cell, independent of cell size (Fig. 5b); we classify this pattern as non-restricted ICM localization. Non-restricted ICM localization was observed for both ICM architectures; while four species utilized vesicular ICMs, the other three utilized lamellar ICMs (Fig. 5a). EM imaging of three of these species illustrated that both architectures indeed spanned along the length of the cells, albeit in slightly different manners. *Rvu. sulfidophilum* contained ICM vesicles throughout the cytoplasm (Fig. 5c), similar to well-studied *Rsp. rubrum*, *Rba. sphaeroides*, and *Rba. capsulatus*^24,36,37^. Meanwhile, both *Rsp. centenum* and *Rbl. acidophilus* harbored ICM lamellae stacks that hugged the cell periphery (Fig. 5c); however, while *Rsp. centenum* ICMs ran around the entire cell, including the pole, *Rbl. acidophilus* ICMs terminated prior to reaching the pole tip, consistent with the absence of autofluorescence at the tips of *Rbl. acidophilus* cells (Fig. 5b). In *Rbl. acidophilus* cells we frequently observed a narrow line of decreased autofluorescence along the longitudinal mid-line of the cell (Fig. 5b), which presumably represented the interior portion of the cytoplasm devoid of ICM. We did not observe a similar line in *Rsp. centenum* or *Rbi. orientis*; however, cells of these two species were narrower and exhibited weaker autofluorescence, and thus autofluorescence likely appeared uniform rather than peripheral due to inadequate resolution.

The six remaining species examined by microscopy all utilized lamellar ICMs and exhibited restricted ICM localization. Five species (*Afifella marina*, *Afifella pfennigii*, *Rhodoplanes piscinae*, *Rhodoplanes elegans*, and *Rhodoplanes roseus*) exhibited autofluorescence patterns resembling that observed in *Rps. palustris* (Fig. 5b). Lending support to this similarity, EM imaging of *Rpl. piscinae* showed peripheral ICM lamellae stacks localized to discrete regions near the cell pole (Fig. 5c). Analysis of *Rhodoplanes* autofluorescence indicated that the pattern of ICM restriction was indistinguishable from that of *Rps. palustris*: ICM number and spacing correlated with cell length, single ICMs were located near the adhesin-producing old pole, and second ICMs maintained a similar distance from the new pole (Fig. 6, Supplementary Fig. 8). We conclude that the same pattern of longitudinal ICM restriction is shared between members of the *Rhodopseudomonas* and *Rhodoplanes* genera, and likely *Afifella* as well. In the sixth species, *Rhodomicrobium vannielii*, autofluorescence varied in both shape and uniformity of intensity between cells but localized away from the region of the cell body from which reproductive filaments emerged (Fig. 5b). Prior EM studies of *Rmi. vannielii* showed irregular ICM density and shape within cell bodies and the absence of ICMs and photosynthetic pigments within filaments^6,38,39^, consistent with the autofluorescence observed herein. Given that ICMs are limited to the cell body and are not continuous between adjoined *Rmi. vannielii* cells, we have chosen to classify *Rmi. vannielii* as having restricted ICM localization, along with the *Afifella* and *Rhodoplanes* species.

**Fig. 6.**
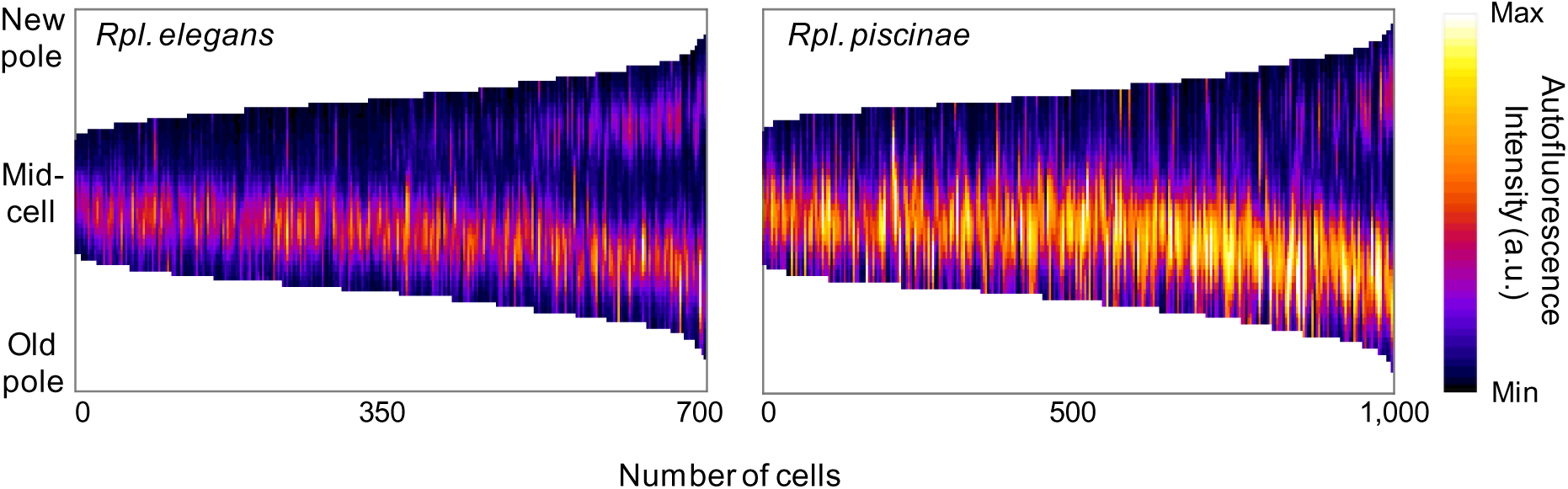
Longitudinally-restricted ICM localization in *Rpl.elegans* and *Rpl. piscinae* cells. Demographs of autofluorescence intensities measured along the medial cell axis of *Rpl. elegans* and *Rpl. piscinae* cells. Cells were sorted from shortest to longest, and polarity was determined by fluorescent lectin staining of the unipolar polysachharide adhesin at the old cell pole. *Rpl. elegans*, n = 714 cells; *Rpl. piscinae*, n = 1001 cells.

While neither ICM localization nor architecture strictly correlated with phylogeny (Fig. 5), we were able to draw several inferences about the evolutionary history of ICM patterning. All members of Rhodospirillales and Rhodobacterales exhibited non-restricted localization, while Rhizobiales members exhibited both restricted and non-restricted localization. We therefore infer an ancestral state of non-restricted ICM patterning, with restricted localization arising as a late innovation in the Rhizobia after their split from the Rhodobacterales (Fig. 5). Given that restricted localization appears in only seven of the nine Rhizobiales species studied here (*Rbi. orientis* and *Rbl. acidophilus* appear to have derived, non-restricted ICM patterns) while the lamellar architecture has been maintained across the order, we further infer that the ancestral PNSB Rhizobiales species had restricted lamellar ICMs. Separately, we infer that the ancestral PNSB Rhodobacterales species had non-restricted vesicular ICMs, as these traits are conserved across the order. With regard to the broader evolutionary history of ICM architecture, Rhodobacterales and Rhizobiales each utilized a single architecture (vesicles and lamellae, respectively), while Rhodospirillales contained members with each architecture type, and these types did not cluster as separate lineages. The presence of both architectures within Rhodospirillales confounds any inference of the ancestral PNSB ICM architecture. Overall, we conclude that ICMs are subject to differential spatial organization among PNSB but not in strict accordance with ICM architecture or species phylogeny.

## DISCUSSION

Here we revealed diversity in ICM localization among PNSB by exploiting the natural autofluorescence of BChl. Because BChl is a native component of the ICM-residing photosynthetic machinery, this method did not require genetic manipulation (i.e. fluorescent protein fusions), thereby enabling characterization of diverse PNSB for which genetic tools are unavailable. However, this approach has limitations. Autofluorescence intensity correlates with cellular BChl content (Fig. 2e), thus PNSB with prohibitively low cellular BChl content will not be amenable to this approach. By the same token, an absence of autofluorescence could be indicative of ICM abundance rather than the absence of ICMs. Thus, absence of autofluorescence should be interpreted carefully and preferably corroborated using additional techniques such as EM, as was done here. Relatedly, the presence of autofluorescence indicates general localization but not ICM architecture. For example, non-restricted autofluorescence was derived from both ICM lamellae and ICM vesicles (Fig. 5). And while restricted localization observed herein was specific to lamellar ICMs in a subset of Rhizobiales members, it is possible that examination of additional species would identify non-Rhizobiales and/or vesicle-bearing PNSB with restricted ICM localization.

We were intrigued by the diversity of ICM localization in the examined PNSB, specifically the differential use of available cell space. Many species elaborated ICMs throughout the cell but other species utilized only specific portions of the cell for ICMs (Fig. 5b), despite that all ICMs presumably serve to enhance light capture and energy transformation^5,8,9^. Our results therefore indicate that ICM function does not necessitate maximizing surface area (i.e. using all available cell space). An ensuing question, then, is whether different localization patterns impart physiological benefits. Perhaps ICM localization diversity reflects differences in the photosynthetic capacity of ICMs and/or the cellular energy demands among PNSB, wherein certain species can transform sufficient energy using only a portion of their cell space and thereby reduce the metabolic costs of ICM synthesis. A non-exclusive possibility is that ICM localization diversity reflects evolution amid other facets of cell physiology, such as cell division. Both non-restricted and restricted ICM localization would be expected to promote ICM inheritance, thereby ensuring that the metabolic (in this case, photosynthetic) potential of cells is maintained across the cell cycle, as is true for protein-encapsulated, CO_2_-fixing carboxysome compartments^40^. However, ICM localization might need to be compatible with the mode of cell growth and division in a given species. While there was no strict correlation between ICM architecture or localization and mode of cell division (binary fission or budding) in the examined PNSB (Supplementary Table 1), there is currently a limited understanding of molecular determinants and mechanisms distinguishing different modes of bacterial growth and division. We anticipate that future insights into bacterial reproduction mechanisms will help clarify any connections between ICMs and cell growth and division. Separately, because ICMs are conditionally made in PNSB^14-16^, ICM localization patterns could differentially impact population fitness during shifts between phototrophic and non-phototrophic (e.g., aerobic) conditions. For example, longitudinally-restricted ICM localization could increase population heterogeneity upon phototrophic-aerobic shift, as mother cells would be expected to retain ICMs while generating ICM-lacking daughter cells. In contrast, PNSB with unrestricted ICMs would be expected to exhibit progressive dilution of ICMs in both mother and daughter cells through rounds of cell division. Additional studies are needed to interrogate the impact of ICM inheritance on cell fitness under both constant phototrophic conditions and upon shifts to aerobic conditions.

Apart from potential benefits conferred by ICM localization, another fundamental question remains: How is ICM localization established and maintained? Presumably, ICM localization is the collective result of the site(s) of ICM synthesis and the extent to which ICMs can expand. In *Rba. sphaeroides* ICM vesicles develop from curved sites in the CM ^41^; little is known about the sites of formation for vesicular or lamellar ICMs in other PNSB. Molecular determinants that influence ICM localization are currently unknown but could include cytoskeletal-like proteins, which are known to coordinate the spatial arrangement of two other bacterial compartments, carboxysomes^40^ and the membrane-bound, magnetic field-sensing magnetosomes^42,43^. PNSB within Rhizobiales lack the actin homolog MreB, but all PNSB have predicted cytoskeletal-like ParA proteins that could be involved in ICM localization. PNSB might also harbor unpredicted cytoskeletal-like proteins, as these proteins are highly divergent and are thus difficult to identify from sequence alone^44^. The diversity in ICM localization revealed herein (Fig. 5) suggests that different synthesis and/or restriction mechanisms are at play, even for architecturally-similar ICMs. Identification of molecular factors governing ICM localization will be invaluable to understand the biological relevance of distinct localization patterns and will foster a broader perspective on subcellular organization in bacteria.

## MATERIALS AND METHODS

### Strains, plasmids, and growth conditions

All species, strains, and plasmids used in this study are listed in Supplementary Table 2. *Rhodobacter sphaeroides* 2.4.1 and *Rhodospirillum rubrum* ATCC 11170 were provided courtesy of Gary Roberts (University of Wisconsin, Madison). *Rhodospirillum centenum* SW and *Rhodobacter capsulatus* SB 1003 were provided courtesy of C. Bauer (Indiana University). *Rhodoblastus acidophilus* DSM 137 was provided courtesy of M. Madigan (Southern Illinois University). Seven strains were purchased from DSMZ (Deutsche Sammlung von Mikroorganismen und Zellkulturen GmbH, Braunschweig, Germany): *Afifella marina* DSM 2698; *Afifella pfennigii* DSM 17143; *Rhodobium orientis* DSM 11290; *Rhodoplanes elegans* DSM 11970; *Rhodoplanes piscinae* DSM 19946; *Rhodoplanes roseus* DSM 5909; and *Rhodovulum sulfidophilum* DSM 1374.

PNSB were grown photoheterotrophically in the following media. *Rhodopseudomonas palustris*, *Rba. sphaeroides*, *Rsp. rubrum*, and *Rhodomicrobium vannielii* were cultivated in defined mineral (PM) medium^45^ supplemented with 5 mM succinate and 0.1% (w/v) yeast extract (YE) (PMsuccYE). *Rsp. centenum* was cultivated on CENS medium^46^. *Rba. capsulatus* was cultivated in modRM2 medium (RM2^47^ modified by substitution of trace elements solution^48^ for trace elements solution 8) supplemented with 5 mM succinate and 0.1% (w/v) YE. All three *Rhodoplanes* species were cultivated in modRM2 supplemented with 20 mM pyruvate and 0.1% (w/v) YE. Marine species were cultivated in modRM2 supplemented with 20 mM pyruvate and 0.1% (w/v) YE, and either 2% (w/v) NaCl (*Afifella* species and *Rvu. sulfidophilum*) or 5% (w/v) NaCl (*Rbi. orientis). Rbl. acidophilus* was cultivated in modABM (acidic basal medium^49^ modified by substitution of trace elements solution^48^ for the trace elements solution of Pfennig and Lippert) supplemented with 20 mM succinate. For photoheterotrophic cultures, media were made anaerobic by bubbling with Ar and then sealing with rubber stoppers and screw caps. Cultures were incubated statically at 30°C at discrete distances from 60 W (750 lumens) soft white halogen bulbs to achieve light intensities as described in the text. Anaerobic, N_2_O-respiring *Rps. palustris* cultures were cultivated in sealed tubes containing anaerobic PM supplemented with 10 mM butyrate, 40 μΜ NaNO3 (necessary for induction of *nos* genes required for N_2_O reduction) (unpublished), and flushed with N_2_O prior to static incubation at 30°C in darkness. Aerobic *Rps. palustris* cultures were grown in 20 mL PMsuccYE in 125 mL flasks at 30°C in darkness with shaking at 225 rpm.

### Generation of *Rps. palustris* mutants

The *crtl* deletion vector (pJQcrtIKO) was generated by PCR amplifying flanking DNA regions using primers pairs BL530/BL531 and BL532/BL533. The two products were fused by *in vitro* ligation at their engineered *sacI* sites. The fusion was subsequently amplified by PCR using primers BL530/BL533 and then ligated into the *xbaI* site of pJQ200SK. The *bchXYZ* deletion vector (pJQbchXYZKO) was generated by PCR amplifying flanking DNA regions using primers pairs BL557/BL558 and BL559/BL560. These primers were designed using the NEBuilder tool (New England Biolabs) for isothermal assembly into plasmid pJQ200SK. PCR products were mixed with xbal-digested pJQ200SK, assembled using Gibson Assembly (NEB), and transformed into NEB10β *E. coli* (New England Biolabs) cultivated on Luria-Bertani medium (Difco).

Deletion vectors were introduced into *Rps. palustris* by electroporation^50^. Mutants were generated using sequential selection and screening as described^51^. Mutant genotypes were confirmed by PCR and sequencing. When necessary, gentamicin was included at 15 μg/mL for *E. coli* or 100 μg/mL for *Rps. palustris*.

### Analytical procedures

Culture growth was monitored by optical density at 660 nm (OD_660_) using a Genesys 20 visible spectrophotometer (Thermo-Fisher). Room temperature absorbance spectra of whole cells resuspended in 60% (w/v) sucrose in PM were recorded using a Synergy MX spectrofluorometer (BioTek). Light intensity was monitored using a LI-250A light meter equipped with a LI-190R quantum sensor (LI-COR).

Bacteriochlorophyll (BChl) *a* was extracted and quantified as previously described^14,52^. Briefly, cells were centrifuged and resuspended in 600 μl phosphate-buffered saline (PBS), at which point sample optical density at 660 nm (OD_660_) was recorded. Cells were then pelleted again, resuspended in 20 μl H_2_O, mixed with 1 ml 7:2 (v:v) acetone:methanol solvent, and incubated at room temperature in darkness for 90 min. Cell debris was removed by centrifugation (max speed, 5 min), and extracted BChl *a* was detected by recording supernatant absorbance at 770 nm (A_770_). BChl *a* content is reported normalized to sample optical density (A_770_/OD_660_).

### Staining polar adhesins with fluorescent lectins

*Rps. palustris* or *Rhodoplanes spp*. cells were resuspended in PBS and incubated with Alexa Fluor 488-conjugated wheat-germ agglutinin (WGA) or concanavalin A (ConA) (Invitrogen), respectively, at a final concentration of 0.5 ug/mL for 5 min prior to fluorescence microscopy imaging.

### Fluorescence microscopy

General cell imaging was performed on 1.5% agarose pads made with unsupplemented media. For timelapse microscopy, log-phase *Rps. palustris* cells from photoheterotrophic cultures grown in 8 μmol s^−1^ m^−2^ light were applied to 1% (w/v) agarose pads made with PMsuccYE and sealed with Valap (1:1:1 vaseline:lanolin:paraffin) prior to imaging. Slides were left incubating on the microscope at room temperature between captures. All imaging was performed on a Nikon Eclipse E800 equipped with a 100x Plan Apo Ph3 DM oil immersion objective, Xcite 120 metal halide lamp (Excelitas Technologies), 83700 DAPI/FITC/Texas Red triple filter cube (Chroma Technologies), and a Photometrics Cascade 1K EM-CCD camera. Images were captured using NIS Elements (Nikon).

### Image analysis

All autofluorescence images were subject to an elastic transformation to correct a consistent spatial distortion resulting from using non-UV-corrected microscope components. A reference transformation matrix was generated from phase contrast and epifluorescence images of a single, crowded field of *Rba. sphaeroides* cells (diffuse autofluorescence) using the bUnwarpJ^53^ plugin in Fiji^54^; the resulting matrix transformation was then applied to images of all 14 species prior to analysis. Detection of cells, autofluorescence, and polar adhesins was performed using MicrobeJ^55^. As necessary, the MicrobeJ Manual Editor was used to split touching cells. All cell shape parameters and autofluorescent foci (ICM) positions were measured by MicrobeJ. Demographs and XY cell maps were generated by MicrobeJ.

### Electron microscopy

Samples were fixed in 2.0% glutaraldehyde, 1% tannic acid in unsupplemented PM medium for one hour at room temperature, with the exception of *Rvu. sulfidophilum* which was fixed in 2.0% glutaraldehyde in modRM2 medium containing 2% (w/v) NaCl. Fixed samples were put on ice and put through four changes of unsupplemented medium, incubation in 1% osmium tetroxide in 0.1 M sodium cacodylate buffer pH 7.2 for one hour, followed by two changes of sodium cacodylate buffer. Samples were then put through a graded ethanol dehydration series to 100% ethanol with *en bloc* staining of 2% uranyl acetate for 30 min at the 75% ethanol step. Samples were subsequently put through three changes of 100% ethanol at room temperature, followed by infiltration using Low Viscosity Embedding resin (Electron Microscopy Sciences, Hatfield, PA) with four changes of resin before the samples were polymerized at 65°C for 18 hours. Ultrathin sections of 85 nm thickness on 300 mesh copper grids were obtained using a Leica Ultracut UCT ultramicrotome. Grids were stained with saturated uranyl acetate and lead citrate, and imaged using a JEM-1400Plus 120 kV transmission electron microscope (JEOL USA, Inc) with a Gatan OneView 4k × 4k camera.

### Genome sequencing

Genome sequencing of the six bacterial strains indicated below was performed at the Indiana University Center for Genomics and Bioinformatics. Paired-end libraries generated with NEXTflex Rapid DNA-seq kit (Bioo Scientific) were sequenced using standard Illumina sequencing protocols. Reads were adapter trimmed and quality filtered using Trimmomatic v0.33, requiring a minimum quality of q20 and a minimum read length of 50 nucleotides after trimming. Reads were assembled using the software SPAdes v3.9.1. Draft genomes were annotated using the NCBI Prokaryotic Genome Annotation Pipeline v4.2 ^56^. These Whole Genome Shotgun projects have been deposited at DDBJ/ENA/GenBank under the following accessions: *Rhodoblastus acidophilus* DSM 137, NPET00000000; *Rhodobium orientis* DSM 11290, NPEV00000000; *Rhodoplanes elegans* DSM 11907, NPEU00000000; *Rhodoplanes piscinae* DSM 19946, NPEW00000000; *Rhodoplanes roseus* DSM 5909, NPEX00000000; *Afifella marina* DSM 2698, NPBC00000000.

### Phylogenetic analysis

Whole genome data were obtained from the genome database at the National Center for Biotechnology Information^57^. Conserved housekeeping gene amino acid sequences were automatically identified, aligned, and concatenated using Phylosift^58^. Maximum likelihood phylogeny reconstruction and bootstrap support estimation were performed in RAxML v8.2.9 ^59^, and trees were visualized using iTOL v4.0.2 ^60^.

## ACKNOWLEDGEMENTS

This work was supported by National Institutes of Health National Research Service Award F32GM112377 to B.L., and National Institutes of Health grant R35GM122556 to Y.V.B. The authors thank members of the McKinlay and Brun labs for discussions, and D. Kearns and A. Randich for comments on the manuscript.

## CONTRIBUTIONS

B.L., D.T.K., B.D.S., Y.V.B., and J.B.M. designed the experiments. B.L., D.T.K., and B.D.S. performed the experiments. A.D. assisted with fluorescence microscopy image analysis. B.L., D.T.K., Y.V.B., and J.B.M. analyzed the data. B.L. and J.B.M. wrote the manuscript. All authors reviewed, edited and approved the manuscript.

## SUPPLEMENTARY INFORMATION

**Supplementary Figure 1.**
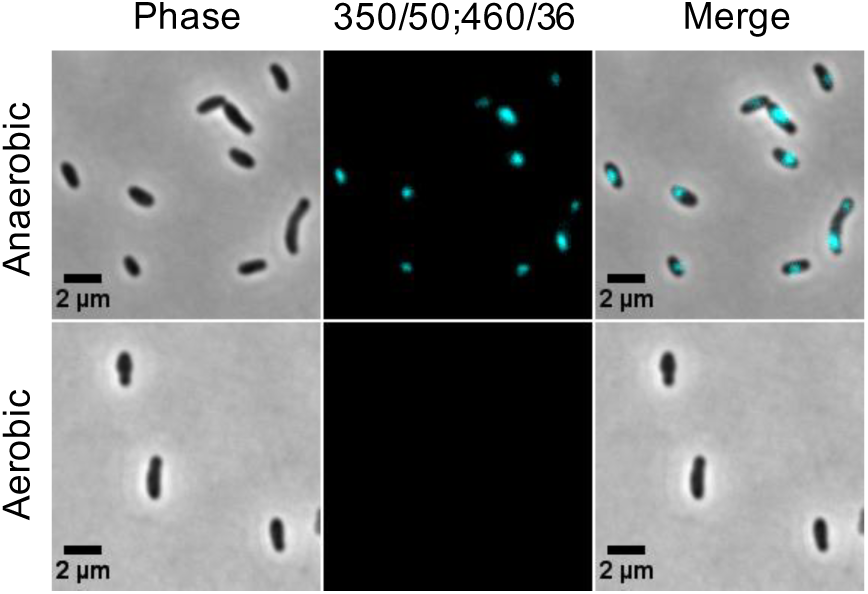
Autofluorescence is specific to ICM-containing *Rps. palustris* cells. Microscopy images of cells grown in PMsuccYE either anaerobically in 8 μmol s^−1^ m^−2^ light or aerobically in darkness, showing ICM-associated autofluorescence visualized using an AT350/50 (DAPI) excitation filter (only the excitation wavelength differs from DAPI filter used in Fig. 2c). LUTs are equivalent between panels.

**Supplementary Figure 2.**
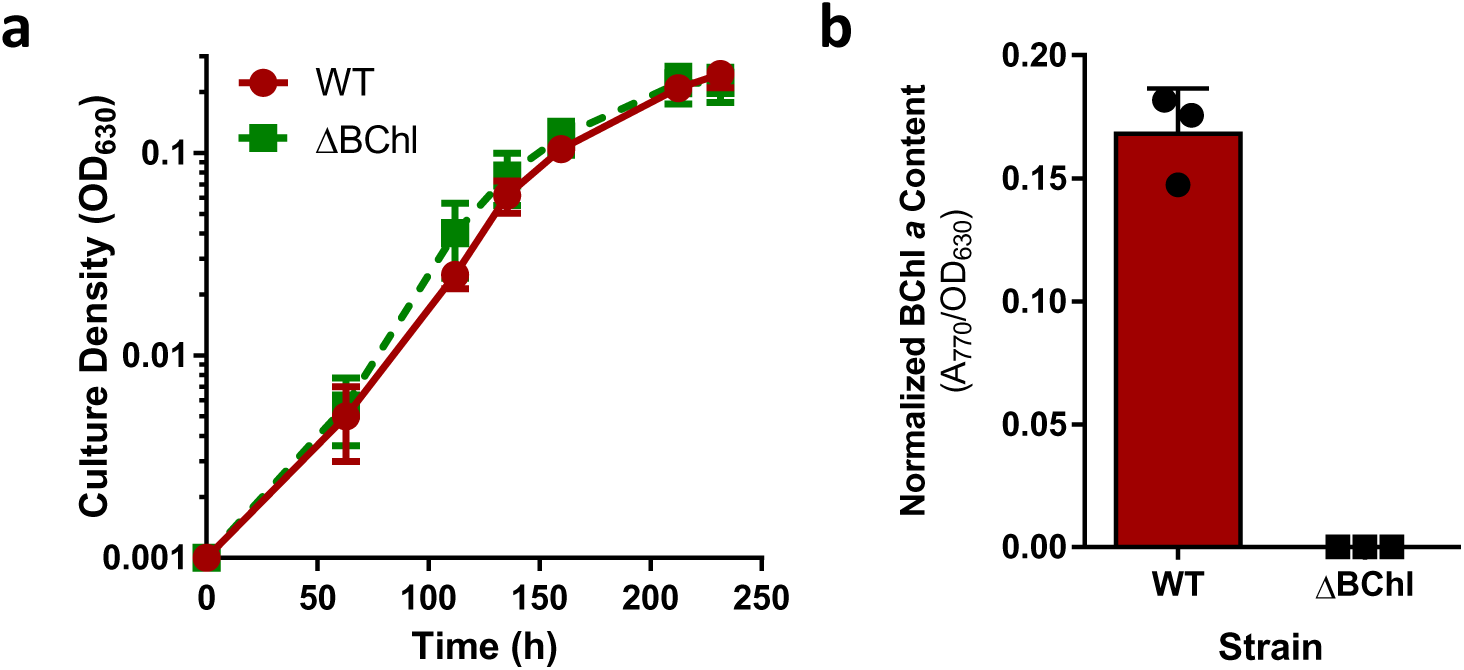
During anaerobic respiration, synthesis of photosynthetic machinery is induced but is dispensable for growth. **a**, Growth curves of WT (dark red circles) and ΔBChl mutant (green squares) *Rps. palustris* grown by anaerobic respiration with N_2_O in darkness. Growth was monitored at OD630 rather than OD_660_ due to the absorbance peak near 660 nm in the ΔBChl mutant (see Fig. 1c). **b**, BChl content of WT and ΔBChl cells, confirming that deletion of *bchXYZ* abolishes BChl synthesis. Error bars, SD, n=3.

**Supplementary Figure 3.**
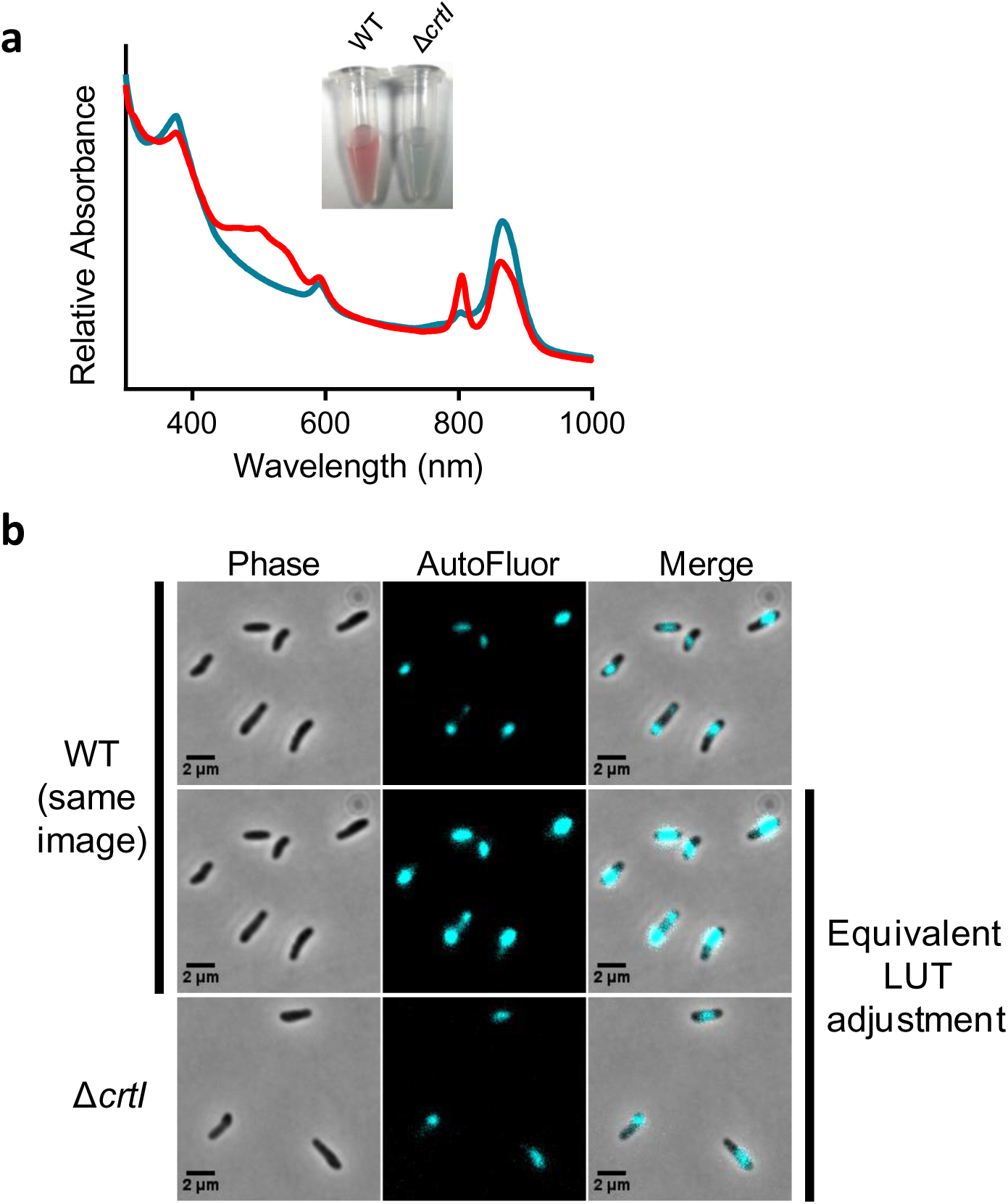
Focal autofluorescence persists in the absence of carotenoids. **a**, Spectral analysis of WT (red) and Δ*crtl* mutant (blue) *Rps.palustris* strains grown in anaerobic PMsuccYE in 20 μmol s^−1^ m^−2^ light, confirming loss of carotenoid synthesis in the mutant. *Inset*, photograph of concentrated cells of each strain. **b**, Microscopy images of WT and Δ*crtl* cells from cultures used in **b**.

**Supplementary Figure 4.**
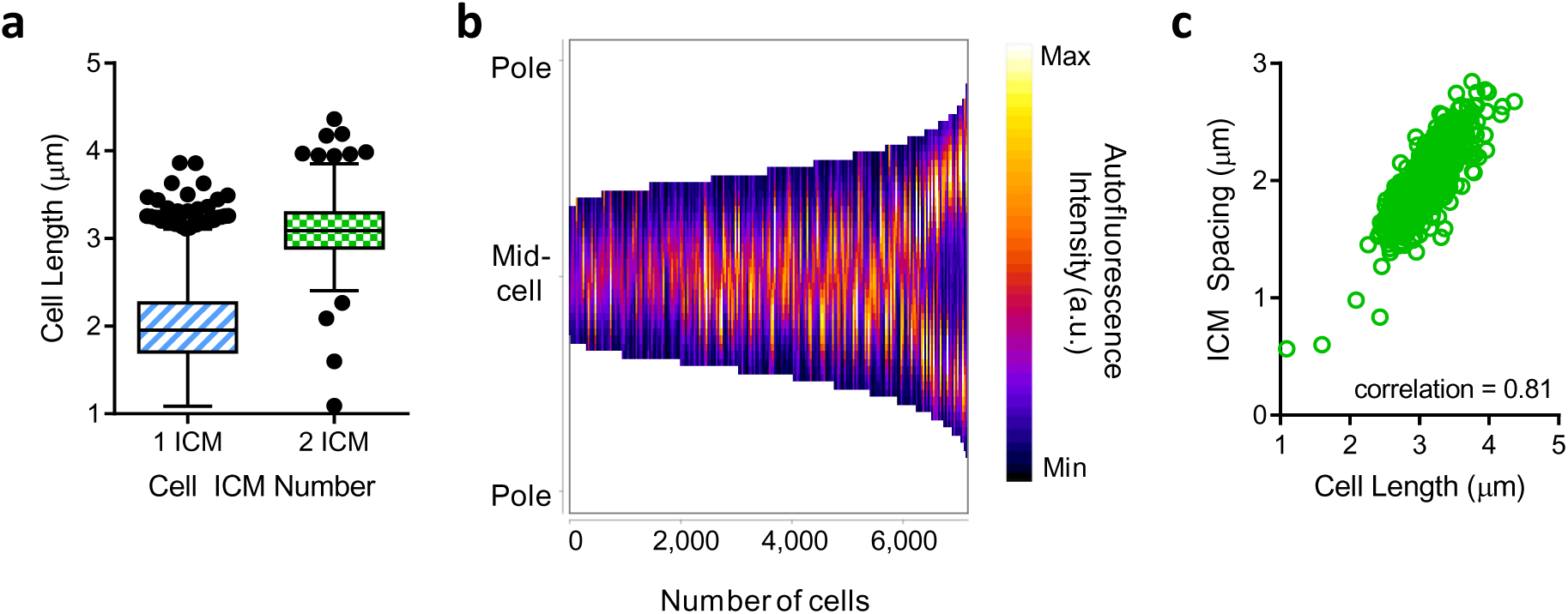
ICM pattern is conserved in non-UPP-bearing *Rps. palustris* cells. **a**, Lengths of cells containing one or two ICMs. 1 ICM, n = 6493 cells; 2 ICM, n = 648 cells. **b**, Demograph of autofluorescence intensities measured along the medial cell axis of all cells in **a**. Cells were sorted from shortest to longest. No cell polarity was assigned. **c**, Longitudinal distance (ICM spacing) between ICMs within cells containing two ICMs plotted as a function of cell length. n = 648 ICM pairs.

**Supplementary Figure 5.**
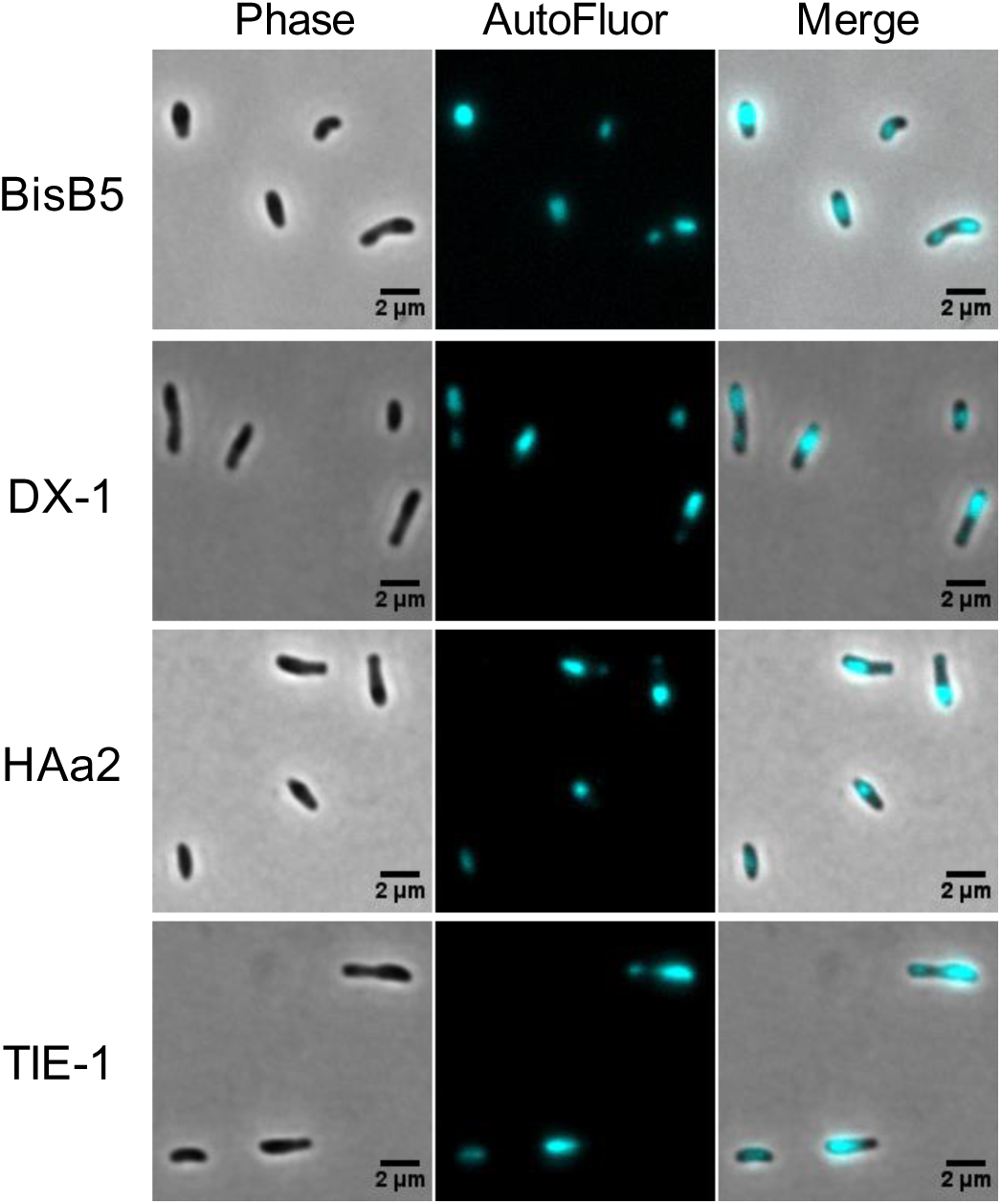
Restricted autofluorescence localization is conserved among diverse ecotypes of *Rps. palustris*. Microscopy images of environmental *Rps. palustris* isolates grown in anaerobic PMsuccYE in 8 μmol s^−1^ m^−2^ light. LUTs are not equivalent between images.

**Supplementary Figure 6.**
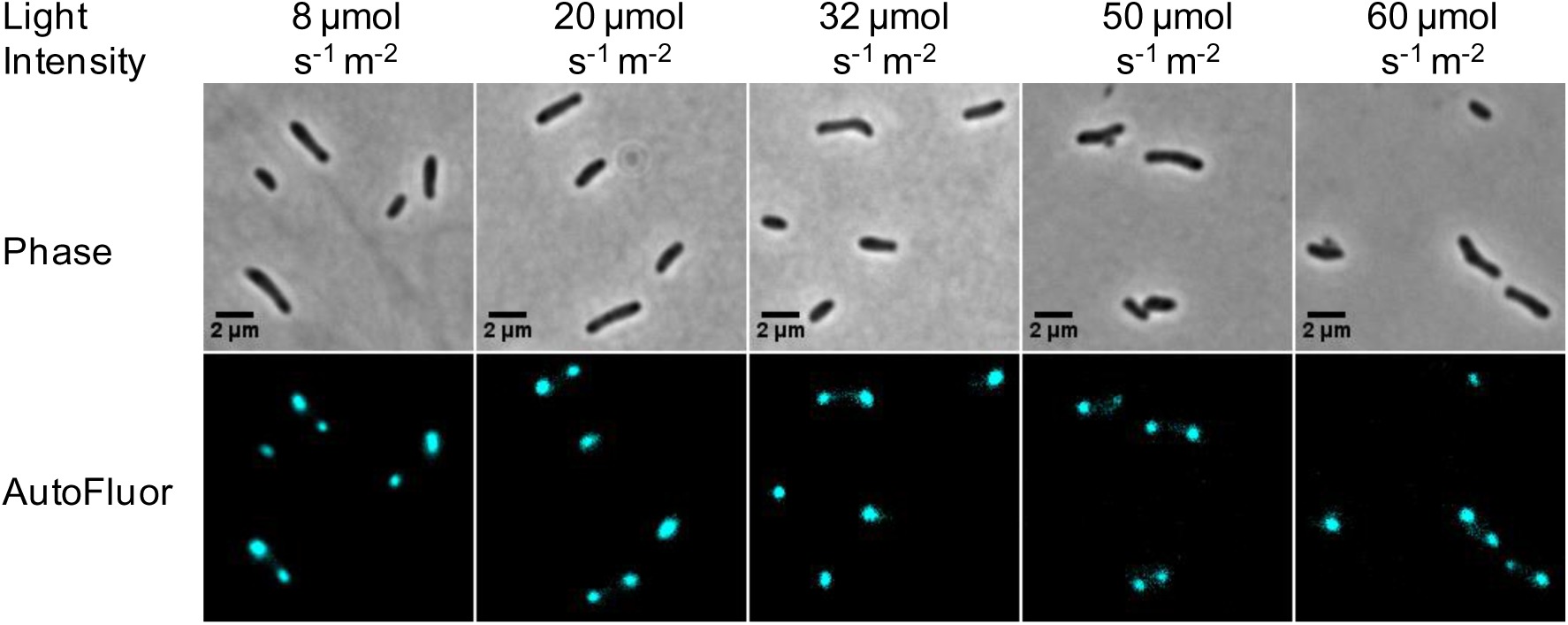
Autofluorescence localization is conserved across a range of light intensities. Microscopy images of *Rps. palustris* CGA009 grown in anaerobic PMsuccYE in different light intensities. Images are from among those used for analysis in Fig. 1d. LUTs are not equivalent between images.

**Supplementary Figure 7.**
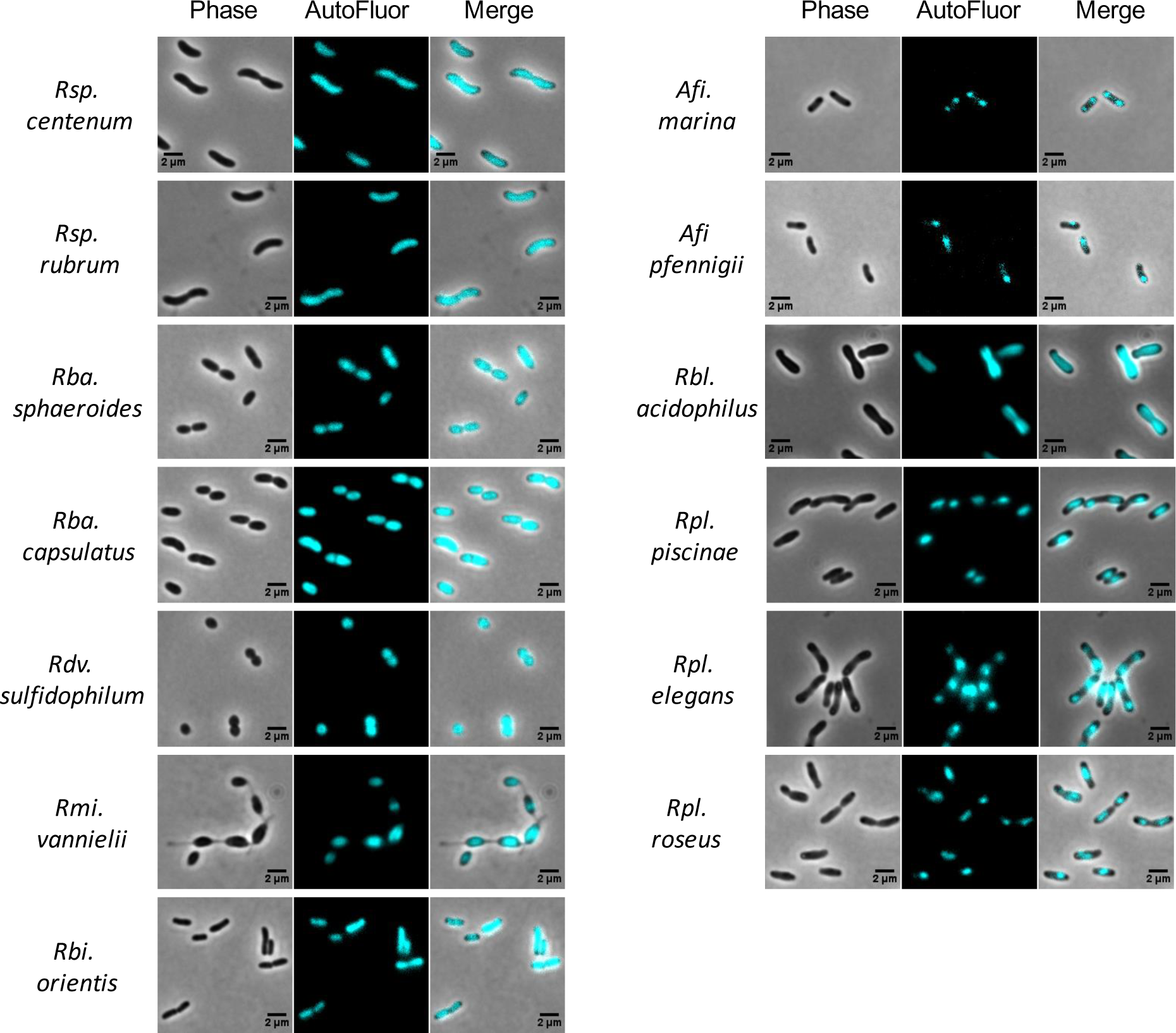
Fluorescence microscopy images of PNSB, with phase and overlay panels included. All species were grown phototrophically in 8 μmol s^−1^ m^−2^ light. LUTs are not equivalent between panels.

**Supplementary Figure 8.**
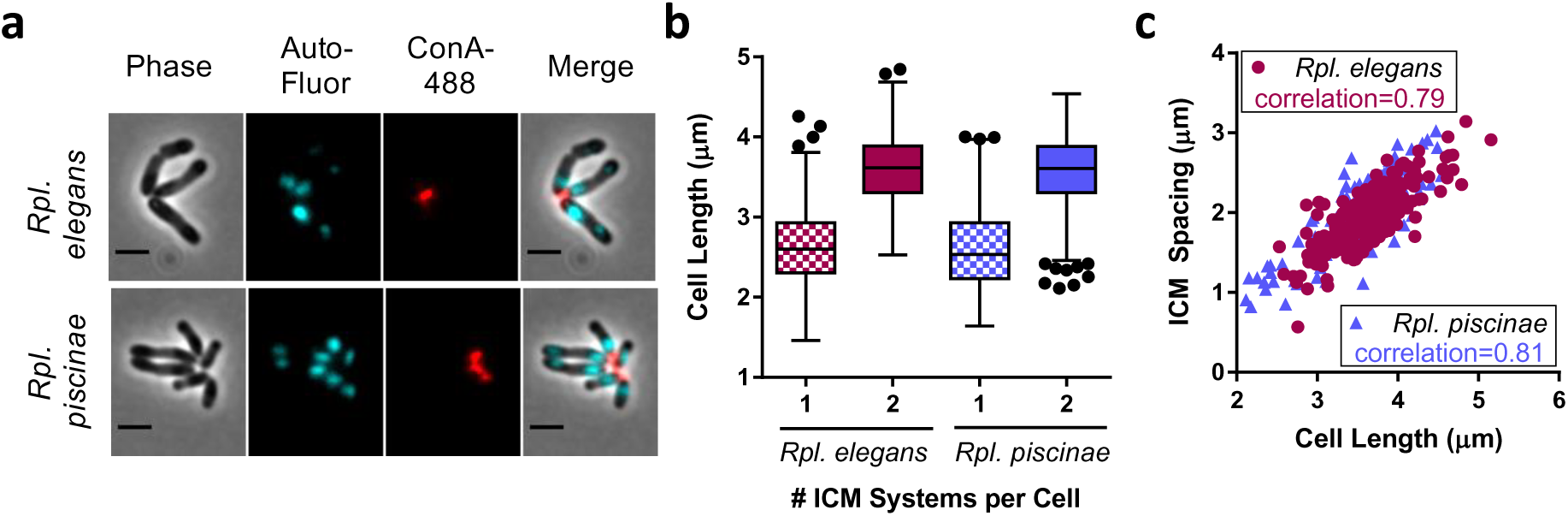
Quantitative analysis of ICM localization in adhesin-bearing *Rpl.elegans* and *Rpl. piscinae* cells. **a**, Microscopy image of cells showing autofluorescence (AutoFluor, cyan) and polar adhesin stained with ConA-488 (false-colored red). Scale bar, 2 μm. **b**, Lengths of cells containing one or two ICMs. *Rpl. elegans*, 1 ICM, n = 496 cells, 2 ICM, n = 218 cells; *Rpl. piscinae*, 1 ICM, n = 846 cells, 2 ICM, n = 155 cells. **c**, Longitudinal distance (ICM spacing) between ICMs within cells containing two ICMs plotted as a function of cell length. *Rpl. elegans*, n = 218 ICM pairs; *Rpl. piscinae*, n = 155 ICM pairs.

**Supplementary Table 1.** Characteristics of select purple non-sulfur bacteria.

| Species<sup>a</sup> | Order | Reproduction Type<sup>b</sup> | ICM Type<sup>c</sup> | BChl Type | Carotenoid Series<sup>d</sup> | Habitat | Reference |
| --- | --- | --- | --- | --- | --- | --- | --- |
| *Rhodovibrio salinarum* | Rhodospirillales | Binary Fission | V | *a* | Spirilloxanthin | Saltern | 1,2 |
| ***Rhodospirillum centenum*** * |  | Binary Fission | L | *a* | Spirilloxanthin | Freshwater | 3-5 |
| *Phaeospirillum molischianum* |  | Binary Fission | L | *a* | Spirilloxanthin | Freshwater | 2,6,7 |
| *Phaeospirillum fulvum* |  | Binary Fission | L | *a* | Spirilloxanthin | Freshwater | 2,7,8 |
| *Roseospirillum parvum* |  | Binary Fission | L | *a* | Spirilloxanthin | Marine | 9 |
| ***Rhodospirillum rubrum*** |  | Binary Fission | V | *a* | Spirilloxanthin | Freshwater | 2,10 |
| *Rhodospira trueperi* |  | Binary Fission | V | *b* | Spirilloxanthin | Marine | 11 |
| ***Rhodobacter sphaeroides*** | Rhodobacterales | Binary Fission | V | *a* | Spheroidene | Freshwater | 12 |
| ***Rhodobacter capsulatus*** |  | Binary Fission | V | *a* | Spheroidene | Freshwater | 12 |
| *Rhodobaca barguzinensis* |  | Binary Fission | V | *a* | Spheroidene | Marine | 13 |
| ***Rhodovulum sulfidophilum*** |  | Binary Fission | V | *a* | Spheroidene | Marine | 14 |
| ***Rhodomicrobium vannielii*** | Rhizobiales | Budding | L | *a* | Spirilloxanthin | Freshwater | 15,16 |
| ***Rhodobium orientis*** |  | Budding | L | *a* | Spirilloxanthin | Marine | 17 |
| ***Afifella marina*** |  | Budding | L | *a* | Spirilloxanthin | Marine | 18,19 |
| ***Afifella pfennigii*** |  | Budding | L | *a* | Spirilloxanthin | Marine | 18,20 |
| ***Rhodoblastus acidophilus*** |  | Budding | L | *a* | Spirilloxanthin | Freshwater | 21,22 |
| *Blastochloris viridis* |  | Budding | L | *b* | Spirilloxanthin | Freshwater | 23,24 |
| ***Rhodoplanes piscinae*** |  | Budding | L | *a* | Spirilloxanthin | Freshwater | 25 |
| ***Rhodoplanes elegans*** |  | Budding | L | *a* | Spirilloxanthin | Freshwater | 26 |
| ***Rhodoplanes roseus*** |  | Budding | L | *a* | Spirilloxanthin | Freshwater | 26,27 |
| ***Rhodopseudomonas palustris*** |  | Budding | L | *a* | Spirilloxanthin | Freshwater | 8,28 |

<sup>a</sup> Species in bold are those empirically examined herein
<sup>b</sup> Modes of reproduction are those historically reported based on phase-contrast microscopy
<sup>c</sup> V, vesicles; L, lamellae
<sup>d</sup> Carotenoid content described in indicated references and/or <sup>29</sup>
\* Species also known as *Rhodocista centenaria*

**Supplementary Table 2.**
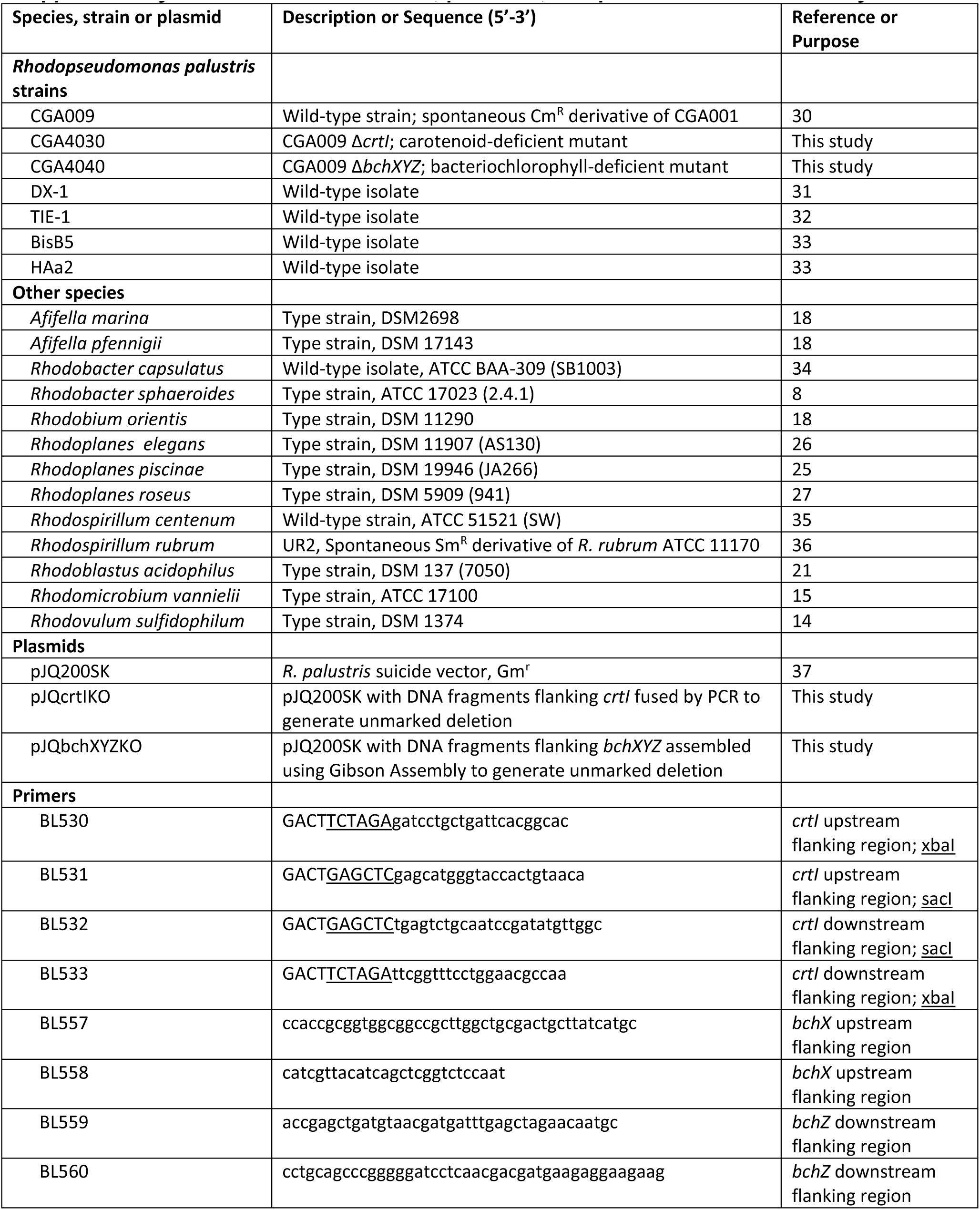
Bacterial strains, plasmids, and primers used in this study.

## REFERENCES

1 Cornejo, E., Abreu, N. & Komeili, A. Compartmentalization and organelle formation in bacteria. Curr Opin Cell Biol 26, 132–138, doi:S0955-0674(13)00194-4[pii]10.1016/j.ceb.2013.12.007 (2014).

2 Madigan, M. T. & Jung, D. O. in The Purple Phototrophic Bacteria eds C. Neil Hunter, Fevzi Daldal, Marion C. Thurnauer, & J. Thomas Beatty) 1–15 (Springer Netherlands, 2009).

3 Drews, G. & Golecki, J. R. in Anoxygenic Photosynthetic Bacteria eds Robert E. Blankenship, Michael T. Madigan, & Carl E. Bauer) 231–257 (Springer Netherlands, 1995).

4 Cohen-Bazire, G. & Kunisawa, R. The fine structure of Rhodospirillum rubrum. J Cell Biol 16, 401–419 (1963).

5 Konorty, M., Kahana, N., Linaroudis, A., Minsky, A. & Medalia, O. Structural analysis of photosynthetic membranes by cryo-electron tomography of intact Rhodopseudomonas viridis cells. J Struct Biol 161, 393–400, doi:S1047-8477(07)00224-9[pii]10.1016/j.jsb.2007.09.014 (2008).

6 Trentini, W. C. & Starr, M. P. Growth and ultrastructure of Rhodomicrobium vannielii as a function of light intensity. J Bacteriol 93, 1699–1704 (1967).

7 Varga, A. R. & Staehelin, L. A. Spatial differentiation in photosynthetic and non-photosynthetic membranes of Rhodopseudomonas palustris. J Bacteriol 154, 1414–1430 (1983).

8 Kis, M., Asztalos, E., Sipka, G. & Maroti, P. Assembly of photosynthetic apparatus in Rhodobacter sphaeroides as revealed by functional assessments at different growth phases and in synchronized and greening cells. Photosynth Res 122, 261–273, doi:10.1007/s11120-014-0026-4 (2014).

9 Kaplan, S., Cain, B. D., Donohue, T. J., Shepherd, W. D. & Yen, G. S. Biosynthesis of the photosynthetic membranes of Rhodopseudomonas sphaeroides. J Cell Biochem 22, 15–29, doi:10.1002/jcb.240220103 (1983).

10 Kiley, P. J., Varga, A. & Kaplan, S. Physiological and structural analysis of light-harvesting mutants of Rhodobacter sphaeroides. J Bacteriol 170, 1103–1115 (1988).

11 Varga, A. R. & Staehelin, L. A. Membrane adhesion in photosynthetic bacterial membranes. Light harvesting complex I (LHI) appears to be the main adhesion factor. Arch Microbiol 141, 290–296 (1985).

12 Siebert, C. A. et al. Molecular architecture of photosynthetic membranes in Rhodobacter sphaeroides: the role of PufX. EMBO J 23, 690–700, doi:10.1038/sj.emboj.76000927600092[pii] (2004).

13 Hsin, J. et al. Self-assembly of photosynthetic membranes. Chemphyschem 11, 1154–1159, doi:10.1002/cphc.200900911 (2010).

14 Cohen-Bazire, G., Sistrom, W. R. & Stanier, R. Y. Kinetic studies of pigment synthesis by non-sulfur purple bacteria. Journal of Cellular and Comparative Physiology 49, 25–68, doi:10.1002/jcp. 1030490104 (1957).

15 Niederman, R. A. Membrane development in purple photosynthetic bacteria in response to alterations in light intensity and oxygen tension. Photosynth Res 116, 333–348, doi:10.1007/s11120-013-9851-0 (2013).

16 Firsow, N. N. & Drews, G. Differentiation of the intracytoplasmic membrane of Rhodopseudomonas palustris induced by variations of oxygen partial pressure or light intensity. Arch Microbiol 115, 299–306 (1977).

17 Sturgis, J. N. & Niederman, R. A. in The Purple Phototrophic Bacteria eds C. Neil Hunter, Fevzi Daldal, Marion C. Thurnauer, & J. Thomas Beatty) 253–273 (Springer Netherlands, 2009).

18 Whittenbury, R. & McLee, A. G. Rhodopseudomonas palustris and Rh. viridis--photosynthetic budding bacteria. Arch Mikrobiol 59, 324–334 (1967).

19 Gunning, B. E. S. & Schwartz, O. M. Confocal microscopy of thylakoid autofluorescence in relation to origin of grana and phylogeny in the green algae. Aust J Plant Physiol 26, 695–708 (1999).

20 Schulze, K., Lopez, D. A., Tillich, U. M. & Frohme, M. A simple viability analysis for unicellular cyanobacteria using a new autofluorescence assay, automated microscopy, and ImageJ. BMC Biotechnol 11, 118, doi:1472-6750-11-118[pii]10.1186/1472-6750-11-118 (2011).

21 Kiley, P. J. & Kaplan, S. Molecular genetics of photosynthetic membrane biosynthesis in Rhodobacter sphaeroides. Microbiol Rev 52, 50–69 (1988).

22 Wraight, C. A., Lueking, D. R., Fraley, R. T. & Kaplan, S. Synthesis of photopigments and electron transport components in synchronous phototrophic cultures of Rhodopseudomonas sphaeroides. J Biol Chem 253, 465–471 (1978).

23 Zeng, X. et al. Proteomic characterization of the Rhodobacter sphaeroides 2.4.1 photosynthetic membrane: identification of new proteins. J Bacteriol 189, 7464–7474, doi:JB.00946-07[pii]10.1128/JB.00946-07 (2007).

24 Holt, S. C. &Marr, A. G. Location of Chlorophyll in Rhodospirillum Rubrum. J Bacteriol 89, 1402–1412 (1965).

25 McEwan, A.G., Greenfield, A. J.,Wetzstein, H. G., Jackson, J. B. & Ferguson, S. J. Nitrous oxide reduction by members of the family Rhodospirillaceae and the nitrous oxide reductase of Rhodopseudomonas capsulata. J Bacteriol 164, 823–830 (1985).

26 Yoshida, K.,Yoshioka, D., Inoue,K., Takaichi, S. & Maeda, I. Evaluation of colors in green mutants isolated from purple bacteria as a host for colorimetric whole-cell biosensors. Appl Microbiol Biotechnol 76, 1043–1050, doi:10.1007/s00253-007-1079-5 (2007).

27 Yildiz, F. H.,Gest, H. & Bauer, C.E. Genetic analys is of photosynthesis in Rhodospirillum centenum. J Bacteriol 173, 4163–4170 (1991).

28 Lang, H. P. & Hunter, C. N. The relationship between carotenoid biosynthesis and the assembly of the light-harvesting LH2 complex in Rhodobacter sphaeroides. Biochem J 298 ( Pt 1), 197–205 (1994).

29 Ng, I. W. et al. Carotenoids are essential for normal levels of dimerisation of the RC-LH1-PufX core complex of Rhodobacter sphaeroides: characterisation of R-26 as a crtB (phytoene synthase) mutant. Biochim Biophys Acta 1807, 1056–1063, doi:S0005-2728(11)00137-X[pii]10.1016/j.bbabio.2011.05.020 (2011).

30 Aagaard, J. & Sistrom, W. R. Control of synthesis of reaction center bacteriochlorophyll in photosynthetic bacteria. Photochem Photobiol 15, 209–225 (1972).

31 Tai, S. P. & Kaplan, S. Intracellular localization of phospholipid transfer activity in Rhodopseudomonas sphaeroides and a possible role in membrane biogenesis. J Bacteriol 164, 181–186 (1985).

32 Bull, M. J. & Lascelles, J. The association of protein synthesis with formation of pigments in some photosynthetic bacteria. Biochem J 87, 15–28 (1963).

33 Whittenbury, R. & Dow, C. S. Morphogenesis and differentiation in Rhodomicrobium vannielii and other budding and prosthecate bacteria. Bacteriol Rev 41, 754–808 (1977).

34 Fritts, R. K., LaSarre, B., Stoner, A. M., Posto, A. L. & McKinlay, J. B. A Rhizobiales-Specific Unipolar Polysaccharide Adhesin Contributes to Rhodopseudomonas palustris Biofilm Formation across Diverse Photoheterotrophic Conditions. Appl Environ Microbiol 83, doi:AEM.03035-16[pii]10.1128/AEM.03035-16 (2017).

35 Oda, Y. et al. Multiple genome sequences reveal adaptations of a phototrophic bacterium to sediment microenvironments. Proc Natl Acad Sci U S A 105, 18543–18548, doi:0809160105[pii]10.1073/pnas.0809160105 (2008).

36 Adams, P. G. & Hunter, C. N. Adaptation of intracytoplasmic membranes to altered light intensity in Rhodobacter sphaeroides. Biochim Biophys Acta 1817, 1616–1627, doi:S0005-2728(12)00170-3[pii]10.1016/j.bbabio.2012.05.013 (2012).

37 Madigan, M. T. & Gest, H. Growth of the photosynthetic bacterium Rhodopseudomonas capsulata chemoautotrophically in darkness with H2 as the energy source. J Bacteriol 137, 524–530 (1979).

38 Morita, S. & Conti, S. F. Localization and Nature of Cytochromes of Rhodomicrobium Vannielii. Arch Biochem Biophys 100, 302–&, doi:Doi 10.1016/0003-9861(63)90077-8 (1963).

39 Conti, S. F. & Hirsch, P. Biology of Budding Bacteria. 3. Fine Structure of Rhodomicrobium and Hyphomicrobium Spp. J Bacteriol 89, 503–512 (1965).

40 Savage, D. F., Afonso, B., Chen, A. H. & Silver, P. A. Spatially ordered dynamics of the bacterial carbon fixation machinery. Science 327, 1258–1261, doi:327/5970/1258[pii]10.1126/science.1186090 (2010).

41 Tucker, J. D. et al. Membrane invagination in Rhodobacter sphaeroides is initiated at curved regions of the cytoplasmic membrane, then forms both budded and fully detached spherical vesicles. Mol Microbiol 76, 833–847, doi:MMI7153[pii]10.1111/j.1365-2958.2010.07153.x (2010).

42 Taoka, A. et al. Tethered Magnets Are the Key to Magnetotaxis: Direct Observations of Magnetospirillum magneticum AMB-1 Show that MamK Distributes Magnetosome Organelles Equally to Daughter Cells. MBio 8, doi:mBio.00679-17[pii]10.1128/mBio.00679-17 (2017).

43 Toro-Nahuelpan, M. et al. Segregation of prokaryotic magnetosomes organellesis driven by treadmilling of a dynamic actin-like MamK filament. Bmc Biol 14, doi:ARTN 88 10.1186/s12915-016-0290-1 (2016).

44 Derman, A. I. et al. Phylogenetic analysis identifies many uncharacterized actin-like proteins (Alps) in bacteria: regulated polymerization, dynamic instability and treadmilling in Alp7A. Mol Microbiol 73, 534–552, doi:MMI6771[pii]10.1111/j.1365-2958.2009.06771.x (2009).

45 Kim, M. K. & Harwood, C. S. Regulation of Benzoate-Coa Ligase in Rhodopseudomonas-Palustris. Fems Microbiol Lett 83, 199–203, doi:Doi 10.1016/0378-1097(91)90354-D (1991).

46 Stadtwalddemchick, R., Turner, F. R. & Gest, H. Physiological-Properties of the Thermotolerant Photosynthetic Bacterium, Rhodospirillum-Centenum. Fems Microbiol Lett 67, 139–143, doi:Doi 10.1016/0378-1097(90)90183-Q (1990).

47 Hiraishi, A. & Kitamura, H. Distribution of Phototrophic Purple Nonsulfur Bacteria in Activated-Sludge Systems and Other Aquatic Environments. B Jpn Soc Sci Fish 50, 1929–1937 (1984).

48 Kremer, T. A., LaSarre, B., Posto, A. L. & McKinlay, J. B. N2 gas is an effective fertilizer for bioethanol production by Zymomonas mobilis. Proc Natl Acad Sci U S A 112, 2222–2226, doi:1420663112[pii]10.1073/pnas.1420663112 (2015).

49 Pfennig, N. Rhodopseudomonas acidophila, sp. n., a new species of the budding purple nonsulfur bacteria. J Bacteriol 99, 597–602 (1969).

50 Pelletier, D. A. et al. A general system for studying protein-protein interactions in Gram-negative bacteria. J Proteome Res 7, 3319–3328, doi:10.1021/pr8001832 (2008).

51 Rey, F. E., Oda,Y. & Harwood, C. S. Regulation of uptake hydrogenase and effects of hydrogen utilization on gene expression in Rhodopseudomonas palustris. J Bacteriol 188, 6143–6152, doi:188/17/6143[pii]10.1128/JB.00381-06 (2006).

52 Schumacher, A.& Drews, G. Effects of light intensity on membrane differentiation in Rhodopseudomonas capsulata. Biochim Biophys Acta 547, 417–428 (1979).

53 Arganda-Carreras, I. et al. in Computer Vision Approaches to Medical Image Analysis: Second International ECCV Workshop, CVAMIA 2006 Graz, Austria, May 12, 2006 Revised Papers eds Reinhard R. Beichel & Milan Sonka) 85–95 (Springer Berlin Heidelberg, 2006).

54 Schindelin, J. et al. Fiji: an open-source platform for biological-image analysis. Nat Methods 9, 676–682, doi:nmeth.2019[pii]10.1038/nmeth.2019 (2012).

55 Ducret, A., Quardokus, E. M. & Brun, Y. V. MicrobeJ, a tool for high throughput bacterial cell detection and quantitative analysis. Nat Microbiol 1, 16077, doi:nmicrobiol201677[pii]10.1038/nmicrobiol.2016.77 (2016).

56 Tatusova, T. et al. NCBI prokaryotic genome annotation pipeline. Nucleic Acids Research 44, 6614–6624, doi:10.1093/nar/gkw569 (2016).

57 Agarwala, R. et al. Database resources of the National Center for Biotechnology Information. Nucleic Acids Research 44, D7–D19, doi:10.1093/nar/gkv1290 (2016).

58 Darling, A. E. et al. PhyloSift: phylogenetic analysis of genomes and metagenomes. PeerJ 2, e243, doi:10.7717/peerj.243243[pii] (2014).

59 Stamatakis, A. RAxML version 8: a tool for phylogenetic analysis and post-analysis of large phylogenies. Bioinformatics 30, 1312–1313, doi:btu033[pii]10.1093/bioinformatics/btu033 (2014).

60 Letunic, I. & Bork, P. Interactive Tree Of Life (iTOL): an online tool for phylogenetic tree display and annotation. Bioinformatics 23, 127–128, doi:btl529[pii]10.1093/bioinformatics/btl529 (2007).

## Supplementary References

1 Nissen, H. & Dundas, I. D. Rhodospirillum-Salinarum Sp-Nov, a Halophilic Photosynthetic Bacterium Isolated from a Portuguese Saltern. Archives of Microbiology 138, 251–256, doi:Doi 10.1007/Bf00402131 (1984).

2 Imhoff, J. F., Petri, R. & Suling, J. Reclassification of species of the spiral-shaped phototrophic purple non-sulfur bacteria of the alpha-Proteobacteria: description of the new genera Phaeospirillum gen. nov., Rhodovibrio gen. nov., Rhodothalassium gen. nov. and Roseospira gen. nov. as well as transfer of Rhodospirillum fulvum to Phaeospirillum fulvum comb. nov., of Rhodospirillum molischianum to Phaeospirillum molischianum comb. nov., of Rhodospirillum salinarum to Rhodovibrio salexigens. Int J Syst Bacteriol 48 Pt 3, 793–798, doi:10.1099/00207713-48-3-793 (1998).

3 Favinger, J., Stadtwald, R. & Gest, H. Rhodospirillum centenum, sp. nov., a thermotolerant cyst-forming anoxygenic photosynthetic bacterium. Antonie Van Leeuwenhoek 55, 291–296 (1989).

4 Yildiz, F. H., Gest, H. & Bauer, C. E. Genetic analysis of photosynthesis in Rhodospirillum centenum. J Bacteriol 173, 4163–4170 (1991).

5 Kawasaki, H., Hoshino, Y., Kuraishi, H. & Yamasato, K. Rhodocista-Centenaria Gen-Nov, Sp-Nov, a Cyst-Forming Anoxygenic Photosynthetic Bacterium and Its Phylogenetic Position in the Proteobacteria Alpha-Group. J Gen Appl Microbiol 38, 541–551, doi:Doi 10.2323/Jgam.38.541 (1992).

6 Giesberger, G. Some observations on the culture, physiology and morphology of some brown-redRhodospirillum-species. Antonie Van Leeuwenhoek 13, 135–148, doi:10.1007/bf02272755 (1947).

7 Takaichi, S., Maoka, T., Sasikala, C., Ramana Ch, V. & Shimada, K. Genus specific unusual carotenoids in purple bacteria, Phaeospirillum and Roseospira: structures and biosyntheses. Curr Microbiol 63, 75–80, doi:10.1007/s00284-011-9941-1 (2011).

8 van Niel, C. B. The Culture, General Physiology, Morphology, and Classification of the Non-Sulfur Purple and Brown Bacteria. Bacteriol Rev 8, 1–118 (1944).

9 Glaeser, J. & Overmann, J. Selective enrichment and characterization of Roseospirillum parvum, gen. nov. and sp. nov., a new purple nonsulfur bacterium with unusual light absorption properties. Arch Microbiol 171, 405–416, doi:91710405.203[pii] (1999).

10 Cohen-Bazire, G. & Kunisawa, R. The fine structure of Rhodospirillum rubrum. J Cell Biol 16, 401–419 (1963).

11 Pfennig, N., Lunsdorf, H., Suling, J. & Imhoff, J. F. Rhodospira trueperi gen. nov., spec. nov., a new phototrophic Proteobacterium of the alpha group. Arch Microbiol 168, 39–45 (1997).

12 Imhoff, J. F., Truper, H. G. & Pfennig, N. Rearrangement of the Species and Genera of the Phototrophic Purple Nonsulfur Bacteria. Int J Syst Bacteriol 34, 340–343 (1984).

13 Boldareva, E. N. et al. Rhodobaca barguzinensis sp. nov., a new alkaliphilic purple nonsulfur bacterium isolated from a soda lake of the Barguzin Valley (Buryat Republic, Eastern Siberia). Microbiology 77, 206–218, doi:10.1134/s0026261708020148 (2008).

14 Hiraishi, A. & Ueda, Y. Intrageneric Structure of the Genus Rhodobacter - Transfer of Rhodobacter-Sulfidophilus and Related Marine Species to the Genus Rhodovulum Gen-Nov. Int J Syst Bacteriol 44, 15–23 (1994).

15 Duchow, E. & Douglas, H. C. Rhodomicrobium Vannielii, a New Photoheterotrophic Bacterium. J Bacteriol 58, 409–416 (1949).

16 Conti, S. F. & Hirsch, P. Biology of Budding Bacteria. 3. Fine Structure of Rhodomicrobium and Hyphomicrobium Spp. J Bacteriol 89, 503–512 (1965).

17 Hiraishi, A., Urata, K. & Satoh, T. A new genus of marine budding phototrophic bacteria, Rhodobium gen. nov., which includes Rhodobium orientis sp. nov. and Rhodobium marinum comb. nov. Int J Syst Bacteriol 45, 226–234, doi:10.1099/00207713-45-2-226 (1995).

18 Urdiain, M. et al. Reclassification of Rhodobium marinum and Rhodobium pfennigii as Afifella marina gen. nov. comb. nov. and Afifella pfennigii comb. nov., a new genus of photoheterotrophic Alphaproteobacteria and emended descriptions of Rhodobium, Rhodobium orientis and Rhodobium gokarnense. Syst Appl Microbiol 31, 339–351, doi:S0723-2020(08)00061-1[pii]10.1016/j.syapm.2008.07.002 (2008).

19 Imhoff, J. F. Rhodopseudomonas-Marina Sp-Nov, a New Marine Phototropic Purple Bacterium. Systematic and Applied Microbiology 4, 512–521 (1983).

20 Caumette, P., Guyoneaud, R., Duran, R., Cravo-Laureau, C. & Matheron, R. Rhodobium pfennigii sp. nov., a phototrophic purple non-sulfur bacterium with unusual bacteriochlorophyll a antennae, isolated from a brackish microbial mat on Rangiroa atoll, French Polynesia. Int J Syst Evol Microbiol 57, 1250–1255, doi:57/6/1250[pii]10.1099/ijs.0.64775-0 (2007).

21 Pfennig, N. Rhodopseudomonas acidophila, sp. n., a new species of the budding purple nonsulfur bacteria. J Bacteriol 99, 597–602 (1969).

22 Imhoff, J. F. Transfer of Rhodopseudomonas acidophila to the new genus Rhodoblastus as Rhodoblastus acidophilus gen. nov., comb. nov. Int J Syst Evol Microbiol 51, 1863–1866, doi:10.1099/00207713-51-5-1863 (2001).

23 Drews, G. & Giesbrecht, P. [Rhodopseudomonas viridis, n. sp., a newly isolated, obligate phototrophic bacterium]. Arch Mikrobiol 53, 255–262 (1966).

24 Hiraishi, A. Transfer of the bacteriochlorophyll b-containing phototrophic bacteria Rhodopseudomonas viridis and Rhodopseudomonas sulfoviridis to the genus Blastochloris gen. nov. Int J Syst Bacteriol 47, 217–219, doi:10.1099/00207713-47-1-217 (1997).

25 Chakravarthy, S. K., Ramaprasad, E. V., Shobha, E., Sasikala, C. & Ramana Ch, V. Rhodoplanes piscinae sp. nov. isolated from pond water. Int J Syst Evol Microbiol 62, 2828–2834, doi:ijs.0.037663-0[pii]10.1099/ijs.0.037663-0 (2012).

26 Hiraishi, A. & Ueda, Y. Rhodoplanes Gen-Nov, a New Genus of Phototrophic Bacteria Including Rhodopseudomonas-Rosea as Rhodoplanes-Roseus Comb-Nov and Rhodoplanes-Elegans Sp-Nov. Int J Syst Bacteriol 44, 665–673 (1994).

27 Janssen, P. H. & Harfoot, C. G. Rhodopseudomonas-Rosea Sp-Nov, a New Purple Nonsulfur Bacterium. Int J Syst Bacteriol 41, 26–30 (1991).

28 Whittenbury, R. & McLee, A. G. Rhodopseudomonas palustris and Rh. viridis--photosynthetic budding bacteria. Arch Mikrobiol 59, 324–334 (1967).

29 Takaichi, S. in The Purple Phototrophic Bacteria eds C. Neil Hunter, Fevzi Daldal, Marion C. Thurnauer, & J. Thomas Beatty) 97–117 (Springer Netherlands, 2009).

30 Kim, M. K. & Harwood, C. S. Regulation of Benzoate-Coa Ligase in Rhodopseudomonas-Palustris. Fems Microbiol Lett 83, 199–203, doi:Doi 10.1016/0378-1097(91)90354-D (1991).

31 Xing, D., Zuo, Y., Cheng, S., Regan, J. M. & Logan, B. E. Electricity generation by Rhodopseudomonas palustris DX-1. Environ Sci Technol 42, 4146–4151 (2008).

32 Jiao, Y., Kappler, A., Croal, L. R. & Newman, D. K. Isolation and characterization of a genetically tractable photoautotrophic Fe(II)-oxidizing bacterium, Rhodopseudomonas palustris strain TIE-1. Appl Environ Microbiol 71, 4487–4496, doi:71/8/4487[pii]10.1128/AEM.71.8.4487-4496.2005 (2005).

33 Oda, Y. et al. Multiple genome sequences reveal adaptations of a phototrophic bacterium to sediment microenvironments. Proc Natl Acad Sci U S A 105, 18543–18548, doi:0809160105[pii]10.1073/pnas.0809160105 (2008).

34 Yen, H. C. & Marrs, B. Map of genes for carotenoid and bacteriochlorophyll biosynthesis in Rhodopseudomonas capsulata. J Bacteriol 126, 619–629 (1976).

35 Lu, Y. K. et al. Metabolic flexibility revealed in the genome of the cyst-forming alpha-1 proteobacterium Rhodospirillum centenum. BMC Genomics 11, 325, doi:1471-2164-11-325[pii]10.1186/1471-2164-11-325 (2010).

36 Singer, S. W., Hirst, M. B. & Ludden, P. W. CO-dependent H2 evolution by Rhodospirillum rubrum: role of CODH:CooF complex. Biochim Biophys Acta 1757, 1582–1591, doi:S0005-2728(06)00311-2[pii]10.1016/j.bbabio.2006.10.003 (2006).

37 Quandt, J. & Hynes, M. F. Versatile suicide vectors which allow direct selection for gene replacement in gram-negative bacteria. Gene 127, 15–21, doi:0378-1119(93)90611-6[pii] (1993).

